# StripePy: fast and robust characterization of architectural stripes

**DOI:** 10.1101/2024.12.20.629789

**Authors:** Andrea Raffo, Roberto Rossini, Jonas Paulsen

## Abstract

**Motivation:** Architectural stripes in Hi-C and related data are crucial for gene regulation, development, and DNA repair. Despite their importance, few tools exist for automatic stripe detection.

**Results:** We introduce StripePy, which leverages computational geometry methods to identify and analyze architectural stripes in contact maps from Chromosome Conformation Capture experiments like Hi-C and Micro-C. StripePy outperforms existing tools, as shown through tests on various datasets and a newly developed simulated benchmark, StripeBench, providing a valuable resource for the community.

**Availability and implementation:** StripePy is released to the public as an open source, MIT-licensed Python application. StripePy source code is hosted on GitHub at https://github.com/paulsengroup/StripePy and is archived on Zenodo. StripePy can be easily installed from source or PyPI using pip and from Bioconda using conda. Containerized versions of StripePy are regularly published on DockerHub.

**Supplementary information:** Supplementary data are provided as a separate file.

## Introduction

Eukaryotic genomes are hierarchically folded inside the nucleus into a 3D conformation crucial for gene regulation, cell division, and DNA repair [5, 6, 14, 61, 65]. At the scale of the nucleus, individual chromosomes form distinct territories [12, 46], further organized into A (euchromatin) and B (heterochromatin) compartments [36]. At the lower levels of the hierarchy, topologically associated domains (TADs) arise from enriched spatial contacts within domains, mediated by CTCF binding at their boundaries [15, 43].

Chromosome conformation capture sequencing methods like Hi-C and Micro-C have been instrumental for revealing these structural hierarchies across the genome. However, deciphering the resulting patterns would be impossible without computational tools designed to detect and analyze them. The significance of these computational methods is underscored by the numerous tools and techniques developed and made available to the research community for analyzing the genome structure at all levels of the genome hierarchy [52] and for generating 3D reconstructions (see, for example, [10, 11]).

In contrast to other genome architectural features, few algorithms are dedicated to the automatic recognition of “ stripes”, which in Hi-C matrices visually consist of vertical (or horizontal) narrow rectangles anchored at the main matrix diagonal. From a biological standpoint, stripes are thought to arise from asymmetric cohesin-mediated loop extrusion [9, 24]. This occurs when loop-extruding cohesin is unidirectionally blocked by a CTCF protein bound to its binding site with the N-terminal end pointing toward it, while loop extrusion can proceed unobstructed on the other side, resulting in a “ stripe” of enriched Hi-C contacts emanating from CTCF bound sites in the genome. Because CTCF binding is enriched at TAD borders, stripes are often found at their edges [9]; this is however not always the case, as it was recently noted that stripes can also appear without a TAD being clearly observed [26].

Stripes are increasingly recognized as important features for epigenetic regulation of transcription and enhancer activity [30, 31], for development [32], and DNA repair [4], highlighting our need to better detect and analyze them. Further, numerous aspects behind loop extrusion are still far from being fully understood, including factors governing the loading of cohesin onto chromatin and the existence of specific targeted loading sites in eukaryotes [16].

Existing stripe detection tools are all rooted in the fields of image processing and analysis. The first method, Zebra [62], identifies pixel tracks of higher interaction frequency at genomic domain boundaries, requiring manual processing to confirm stripe candidates. Zebra’s original implementation is not publicly available, but a re-implementation was made available by an independent group on GitHub under the name of StripeCaller^1^; StripeCaller constrains the stripe width to a single bin and does not assume that stripes are anchored to the main diagonal of the matrix. Another tool published in late 2018 is domainClassifyR^2^. The tool detects architectural stripes based on TAD boundaries by defining a stripe score that is based on the *Z−*statistic; intra-TAD segments remain however undetected. Chromosight [41] exploits template-based pattern recognition by convolving a set of templates over the contact map. A set of criteria is then applied to filter candidate stripes to deal with, e.g., candidate stripes overlapping with too many empty pixels or that are too close to another detected pattern. Chromosight identifies stripes as a single point, without estimating stripe width and height. Stripenn [59] pre-processes the input Hi-C matrix by performing contrast adjustment followed by noise reduction via the Canny edge detection algorithm. A set of custom criteria is introduced to detect and possibly merge vertical lines. Lastly, the tool computes two coefficients – median *p*-value and stripiness – to quantitatively evaluate architectural stripes. Of the four methods available at present, Stripenn is the only one capable of estimating both the width and the height of a stripe, effectively transitioning stripes from a 0- or 1-dimensional pattern to a fully realized 2-dimensional representation.

We present StripePy, a stripe recognition method based on concepts rooted in geometric pattern recognition, topological data analysis [7] and simple geometric reasoning. Implemented in Python, StripePy is uniquely capable of reading interactions in both Cooler (.cool and .mcool) and Juicer (.hic) data formats. Beyond the detection of stripes, including their height and width estimates, StripePy provides a collection of descriptors that can be used for subsequent postprocessing and in-depth analyses. One of the computed descriptors is the *relative change*, which is used to discern candidate stripes that exhibit minimal variation when comparing the average signal within stripes with the average signal in their neighborhood. StripePy thus offers an efficient and user-friendly method to detect architectural stripes, while at the same time computing several descriptors that can be used to inform further downstream analyses.

To assess StripePy’s performance and compare with existing tools, we developed StripeBench, a novel benchmark consisting of 64 Hi-C contact maps simulated with the computational tool MoDLE [56] at different resolutions, contact densities, and noise levels. These maps come with ground truth annotations. Furthermore, StripeBench defines a set of measures to quantify the performance and compare computational tools in the classification of genomic bins and recognition of stripes. StripeBench is used to compare StripePy against StripeCaller, Chromosight, and Stripenn and assess how these methods behave at increasing levels of resolution, contact density, and noise level.

Finally, the analysis of real contact maps from two cells lines is presented, and the predicted stripes are compared against CTCF chromatin immunoprecipitation sequencing (ChIP-Seq) peaks. These analyses demonstrate that StripePy can increase the number of correctly predicted stripes and detected anchor sites while maintaining high overall evaluation scores, making StripePy a valuable contribution to the community.

## Material and methods

### Overview of StripePy

StripePy combines geometric pattern recognition, topological persistence, and simple geometric reasoning to detect stripes in contact maps. StripePy’s CLI consists of three subcommands: `call`, `view`, and `plot`. `stripepy call` is responsible for running the stripe detection algorithm on a contact matrix in Cooler (.cool or .mcool) or Juicer (.hic) format [1, 18] at a given resolution. While StripePy can process both balanced and unbalanced contact maps, tests detailed in the Supplementary Information reveal that our algorithm performs optimally when no prior balancing is applied. Our method depends on a set of parameters, with default values specified both in the remainder of this section and in the help message displayed when executing `stripepy call --help`. The command `stripepy call` produces a file in Hierarchical Data Format (.hdf5) [23] containing a list of rectangular regions corresponding to the candidate stripes together with a set of descriptors and complementary information. The genomic coordinates of the candidate stripes identified by `stripepy call` can be extracted from the .hdf5 file using `stripepy view`, which outputs the stripes coordinates in BEDPE format directly to standard output. Finally, `stripepy plot` can be used to visualize architectural stripes overlaid on top of the Hi-C matrix. `stripepy plot` can also generate several graphs showing the general properties of the called stripes. The remainder of this section summarizes the main concepts behind the StripePy algorithm, which are displayed in the form of the graphically-simplified pipeline in Fig. 1; the reader is referred to the Supplementary Information for the theoretical and technical details, and to Supplementary Fig. 7 for an extended overview of StripePy.

**Fig. 1.**
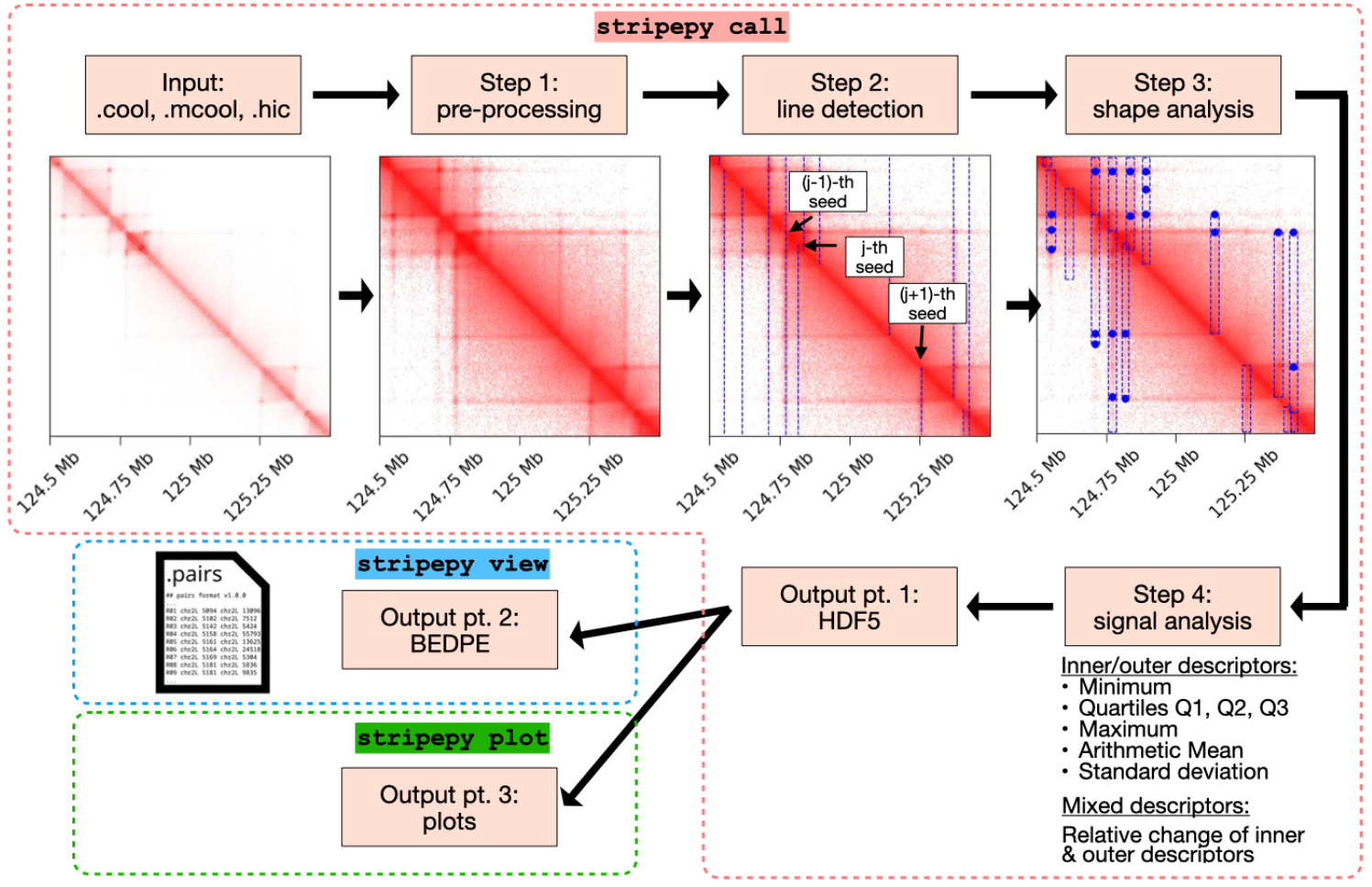
Short overview of StripePy. In `stripepy call`, the input contact map is first pre-processed (Step 1), then studied to detect linear patterns (Step 2) which are subsequently enriched by determining the width and height of each stripe through geometric reasoning, as well as peaks (represented by blue dots) in the signal inside of the stripes (Step 3); the candidates stripes are then post-processed to remove weak occurrences (Step 4). `stripepy view` extracts the genomic coordinates of the candidate stripes and outputs them into a .bedpe file. `stripepy plot` can be used to produce various plots of the steps adopted by `stripepy call`.

StripePy’s algorithm boils down to four consecutive steps:

- In Step 1, StripePy starts by applying a pre-processing step: it consists of a log-transformation, followed by the rescaling of the matrix entries such that they are in [0, 1], and then extracts a band around the main diagonal (here set to the default value of 5 Mbp, which was selected to upper bound TAD sizes in mammals [27]). Then, StripePy searches for vertical stripes in the lower- and upper-triangular parts of the matrix separately by applying the remaining steps independently to the corresponding sub-matrices.
- In Step 2, each of the two triangular matrices is marginalized by summing over the rows to obtain a scalar function of the columns. To mitigate uninformative fluctuations due to columns/rows with low number of interactions, a constraint using the maximum between the scalar function and its weighted moving average is applied [49, 50, 51]. The obtained profile is then scaled so that it has values in [0, 1]. All local maxima are filtered with topological persistence, a technique in topological data analysis requiring a local maximum to stand significantly higher than its surroundings to be considered “ persistent” : said otherwise, only local maxima that are markedly higher from nearby values are retained, while those corresponding to minor fluctuations are disregarded. This allows to keep loci (here called *seed sites*) that exhibit a more marked linear pattern, in a manner similar to the pattern recognition method known as the Hough transform [28, 42]. The default threshold for topological persistence is set to 0.04, i.e., 4%. This value was selected empirically after testing with contact maps generated by MoDLE [56]. The reader is referred to the Supplementary Information for an extended description of how topological persistence works.
- In Step 3, each seed site is analyzed independently to estimate its *horizontal* and *vertical domains* – which consist of the genomic intervals defining the stripe horizontally and vertically, respectively. These can then be used, for example, to compute the stripe width and height and to extract the corresponding regions within the contact map. Given a seed site, the horizontal domain is defined by looking in the neighborhood of the local maximum point where the scalar function from Step 2 is monotonically increasing (resp. decreasing) to the left (resp. to the right) of the maximum point, and then by finding the left (resp. right) bin where the increase (resp. decrease) of the scalar function is the steepest; to prevent excessive stretching of the horizontal domain at low resolutions, a maximum stripe width is introduced (default value: 100 kb). The vertical domain of a stripe is obtained by extracting the columns corresponding to the horizontal domain, marginalizing in a similar manner as in Step 2, and then studying this scalar function. Two criteria are provided: (1) applying topological persistence on the rescaled profile to identify persistent peaks in the signal and use the location of the furthest peak as a boundary; the remaining peaks, if any, point at regions of the stripe where the signal sharply increases; (2) thresholding to a minimum value.
- To enrich what is now just a purely geometric structure (seed site + horizontal domain + vertical domain), Step 4 computes a number of complementary descriptors from the contact map, considering the matrix entries either inside the stripe or outside the stripe (in what we call a *k-neighbour*): minimum, quartiles, maximum, arithmetic mean and standard deviation. We then combine the inner and outer means into the *relative change* parameter, which quantifies to what extent the signal differs between the inside and the outside of the stripe: it is defined as the difference between mean inner and mean outer signals, divided by the mean outer signal; the mean outer signal is computed in the *k-neighbour*.

The .hdf5 file generated from these four steps can be inspected and post-processed using the stripepy view command. Thresholding the relative change equals to ruling out weaker stripes. After conducting tests on datasets from the same cell lines analysed in the manuscript, we have chosen to adopt a threshold of 4% for synthetic data and 3% for real contact maps, which resulted in high single-value scores. Changes in protocols, restriction enzymes, and cell lines may necessitate adjustments of this parameter: lowering the threshold in relative change will retain a greater number of stripes, whereas higher thresholds will lead to more restrictive filtration.

### StripeBench: unified benchmarking of stripe identification algorithms for Hi-C data

A critical challenge in evaluating the performance of any pattern recognition method for Hi-C is the absence of a robust benchmarking system. Despite the wide and ready availability of contact maps at different conditions such as sequencing depth, resolution, and noise level, the absence of controlled experiments, scalability, and standardization hinders the quantitative analysis of – as well as the fair comparison between – existing tools. To overcome these limitations we introduce StripeBench, a benchmark that includes: a set of 64 simulated contact maps containing realistic TAD and stripe patterns, a standardized “ ground truth” baseline for testing, and a collection of metrics for evaluation and comparison. The remainder of this section presents key points and notations related to the benchmark, with a more detailed explanation provided in the Supplementary Information.

The synthetic contact maps are obtained through a genome-wide run of MoDLE [56], a computational tool that uses fast stochastic simulation to sample DNA-DNA contacts generated by loop extrusion. Here, loop extrusion is modelled using extrusion barriers whose occupancy is based on RAD21 ChIP-Seq data from the H1-hESC cell line. Simulations are run using default settings except for the following three factors: target contact density (i.e., the sequencing depth), scale parameter (which controls the noise), and resolution. The effect of changing these parameters on the contact maps is illustrated in Fig. 2A. The ground truth consists of the extrusion barriers (here, CTCF binding sites) inputted to MoDLE. Each barrier includes a position, a blocking direction, and an occupancy value. These three parameters concur to determine the location, direction, and the strength of the linear pattern observed in the resulting contact maps. The ground truth contains 31,423 barriers at 5 kb, with nearly an equal distribution between lower and upper-triangular occurrences. This number decreases to 30,581, 28,631, and 26,191 at 10 kb, 25 kb and 50 kb, respectively: this reduction occurs because we discard duplicated barriers that overlap with the same genomic bin.

**Fig. 2.**
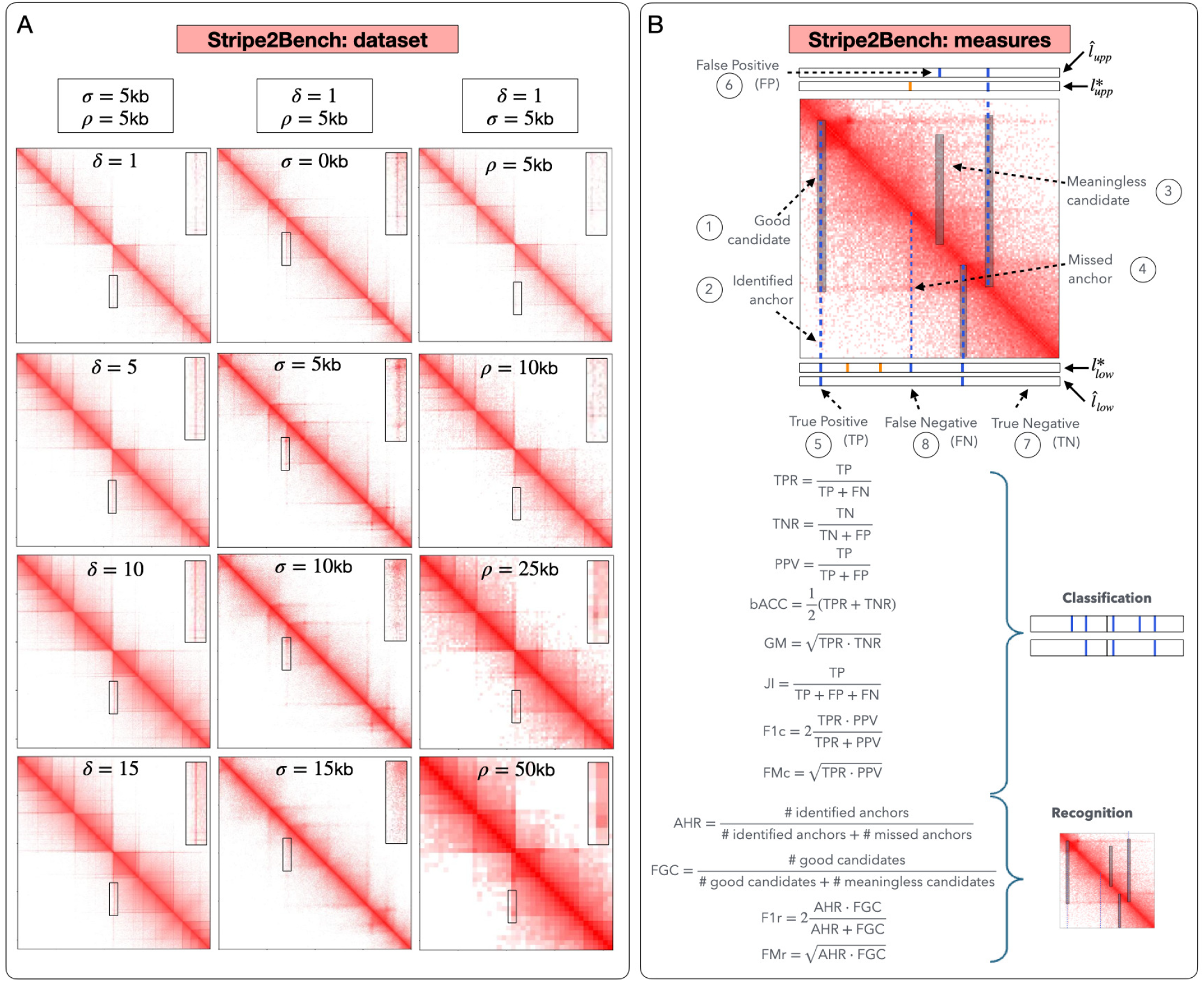
StripeBench. **A** The dataset is composed of 64 contact maps which are generated by varying three factors: contact density *δ*, noise level *σ*, resolution *ρ*; the three columns show the effect of increasing each of these three factors singularly. **B** StripeBench’s recognition vs. classification measures. Circled numbers 1-4 point at concepts relevant in recognition: 1 − 2) show a predicted stripe (semitransparent rectangle) that can be deemed “ good”, as it contains a ground truth anchor point (here represented by a dashed blue vertical line); 3) point at a predicted stripe that is not “ meaningful” as it does not contain any ground truth anchor point; 4) shows a ground truth anchor point that is not found, as it is not contained in any predicted stripe. Circled numbers 5-8 refer, respectively, to an example of true positive, false positive, true negative and false negative in (bin) classification, where we have represented the lower-triangular ground truth classification vector 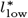 and the predicted classification vectors 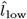 with two rectangles where white means 0 (no anchor point) and vertical colored segments denote 1 (anchor point); blue (resp. orange) refers to anchor points with average extrusion barrier occupancy greater or equal (resp. lower) than 0.70. The panel also lists classification and recognition measures.

To evaluate and compare stripe callers from multiple directions we introduced two families of measures: *classification measures* target the identification of genomic bins hosting extrusion barriers; *recognition measures* finds out whether predicted stripes include bins *hosting* extrusion barriers, i.e., if it is traversed by it. Fig. 2B illustrates these concepts, see also [34].

We adopt the following “ base measures” for the classification problem: sensitivity (TPR), specificity (TNR), and Positive Predictive Value (PPV). Similarly, we introduce the following base recognition measures: *Anchor Hit Rate* (AHR) and *Fraction of Good Candidates* (FGC), which correspond to TPR and PPV in the context of recognition. Despite their popularity and ease of interpretability, these metrics run the risk of being severely biased towards the majority class when applied to highly imbalanced datasets. As a result, they can be misleading if used in isolation [2, 25, 39]. To address these concerns and capture a more balanced evaluation, the following single-value metrics are adopted: balanced accuracy (bACC) and Geometric Mean (GM), which combine sensitivity and specificity, respectively; F1-score (F1c) and Fowlkes–Mallows index (FMc), which summarize precision and recall; Jaccard Index (JI).

## Results

### StripePy: improved stripe classification and recognition

We applied StripePy, Chromosight, StripeCaller, and Stripenn – the currently available stripe callers – to the 64 contacts maps in StripeBench, obtaining a number of predictions that varies with resolution, contact density, and noise level. Specifically, we obtained: from 7,705 to 35,068 stripes with StripePy, from 3,500 to 34,912 with Chromosight, from 0 to 33,002 with StripeCaller, and from 0 to 3,092 with Stripenn. The number of predicted stripes for each contact map is provided in Supplementary Table 1.

Some representative examples of predicted stripes are provided in Fig. 3A. As mentioned before, Chromosight does not provide estimates for the length or width of a stripe; instead, it returns two genomic pairs – each of which is positioned at a distance determined by the chosen resolution. StripeCaller provides an estimate for the length of a stripe, but, similarly to Chromosight, it sets the width to equal the matrix resolution.

**Fig. 3.**
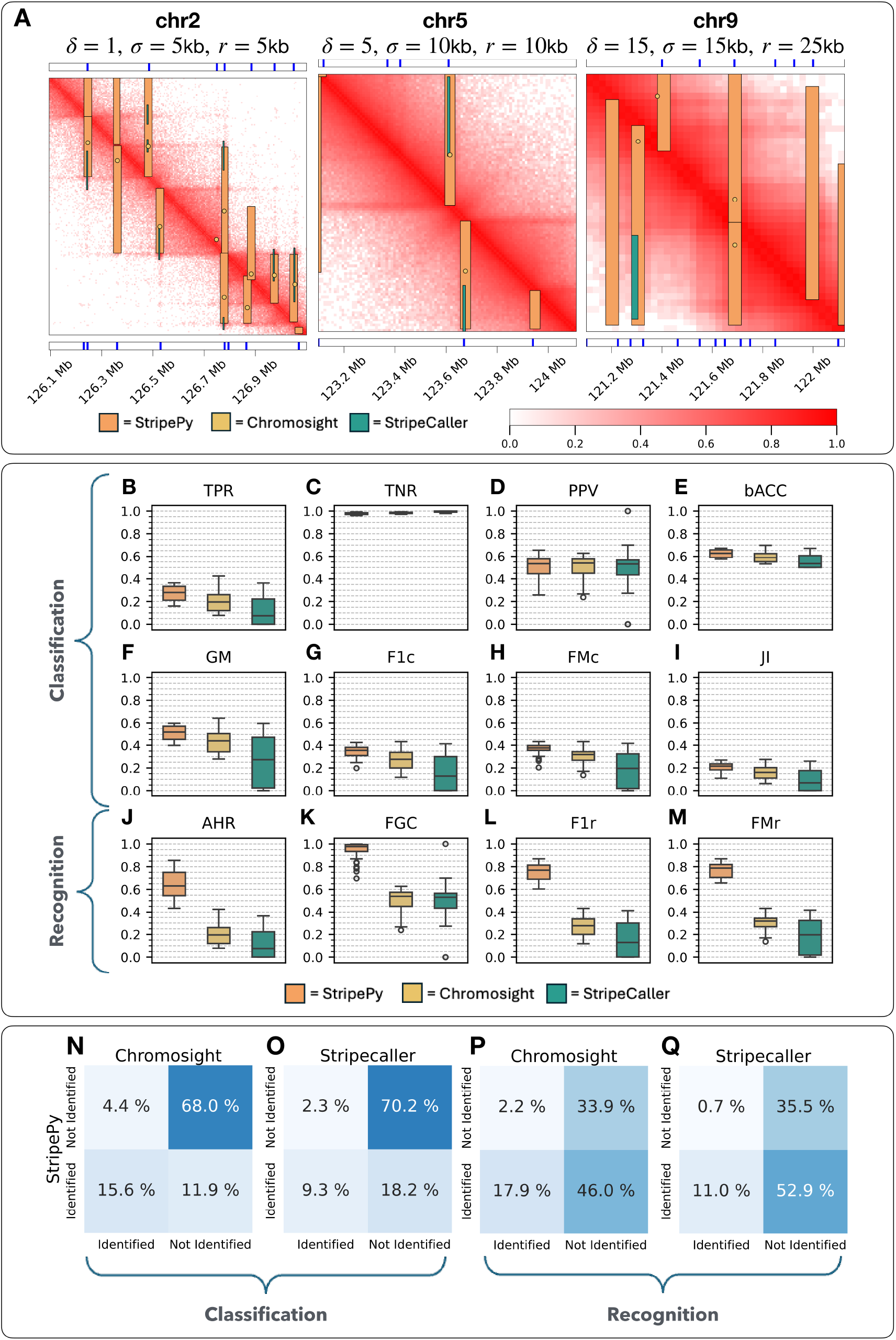
Benchmarking pt. 1. **A** Three regions extracted from three MoDLE-simulated contact maps. The upper and lower bands show the anchor points from the ground truth as solid vertical blue lines, and they refer to the upper- and the lower-triangular parts, respectively. The images of the contact maps have superimposed the predictions produced by the tools under study. **B**-**M** Boxplots of the classification (TPR, TNR, PPV, bACC, GM, F1c, FMc, and JI) and recognition (AHR, FGC, F1r, and FMr) measures over the StripeBench contact maps. **N**-**Q** Heatmaps summarizing how StripePy performs compared to Chromosight and StripeCaller in terms of percentages of ground truth anchor points: (classification) correctly labeled as such; (recognition) found inside a candidate stripe.

Due to the low predictive capability of Stripenn on this dataset, we decided to exclude this tool from the comparative analysis. Nevertheless, its classification and recognition scores on each contact map are included, together with the scores of the other stripe callers, in Supplementary Tables 2-5.

#### StripePy improves overall bin classification

First, we conducted an overall analysis by evaluating each tool’s classification performance on anchor and non-anchor sites using base measures, considering the entire dataset. Results indicate that StripePy markedly outperforms the other tools in the True Positive Rate (TPR) with a median TPR of 0.2794, compared to Chromosight (0.1957) and StripeCaller (0.0753), see Fig. 3B. Most tools show high True Negative Rate (TNR) due to the overwhelming number of non-anchor bins (median TNR values for StripePy, Chromosight, StripeCaller are 0.9756, 0.9833 and 0.9949, respectively, as shown in Fig. 3C). The strong skewness in label distribution has also been shown to affect negatively Positive Predictive Value (PPV), as pointed out in [2]: the median values for StripePy, Chromosight, and StripeCaller are 0.5341, 0.5395, and 0.5314, respectively, see Fig. 3D).

To address the imbalance in positive and negative bins, we selected a weighted accuracy measure (bACC), which balances sensitivity and specificity through an arithmetic mean. StripePy again outperforms the other tools with a median bACC of 0.6272, compared to Chromosight (0.5886) and StripeCaller (0.5349), see Fig. 3E. Applying the geometric mean, which combines TPR and TNR in an alternative fashion, also confirms this trend: median values for StripePy, Chromosight, and StripeCaller are 0.5208, 0.4389, and 0.2736, respectively (Fig. 3F). Examining the harmonic mean of precision and recall (F1c) reinforces StripePy’s improved predictive performance with a median F1c of 0.3554, compared to Chromosight (0.2767) and StripeCaller (0.1291), see Fig. 3G. The alternative combination of precision and recall via the Fowlkes-Mallows index (FMc) also shows StripePy’s superiority with a median FMc of 0.3765, versus Chromosight (0.3181) and StripeCaller (0.1966), see Fig. 3H.

Using the Jaccard Index (JI), a summary of classifier performance that considers true positives, false positives, and false negatives, shows that StripePy outperforms the other tools with a median JI of 0.2161, compared to 0.1605 for Chromosight and 0.0690 for StripeCaller, see Fig. 3I.

Regarding interquartile ranges, StripePy provides the lowest values for all classification measures except for TNR (StripePy: 0.0148, Chromosight: 0.0093, StripeCaller: 0.0090, see Fig. 3C) and PPV (StripePy: 0.1307, Chromosight: 0.1252, StripeCaller: 0.1315, see Fig. 3D. The presence of outliers points to possible limit cases where performance drops significantly (see, for example, the case of TPR in Fig. 3B).

To conclude, StripePy consistently outperforms Chromosight and StripeCaller across various classification metrics, including the base measure True Positive Rate (TPR) and all single-value measures: balanced accuracy (bACC), Geometric Mean (GM), F1 score (F1c), Fowlkes-Mallows index (FMc), and Jaccard Index Coefficient (JI). Chromosight and StripeCaller call fewer stripes with a slightly higher Positive Predictive Value (PPV). In terms of stability, interquartile ranges indicate a generally more stable performance for StripePy. However, unhandled edge cases may still occur, as shown by the presence of outliers.

#### Inclusion of stripe width estimation further increases StripePy’s performance

To understand the impact of width estimation, we conducted a similar analysis focused on the recognition problem. The base measure Anchor Hit Rate (AHR), which corresponds to TPR in this task, confirms that StripePy significantly increases the fraction of recognized anchor sites when estimating widths. The median AHR reaches 0.6312, compared to the median classification measure of 0.2794 (Fig. 3J). Analogously, width estimation also enhances the fraction of positives that are true positives: while StripePy’s median PPV was 0.5341, its counterpart in the recognition task (Fraction of Good Candidates, FGC) soars to 0.9767 (Fig. 3K). As a result, the single-value measures see a substantial rise: the harmonic mean of AHR and FGC (F1r) attains a median value of 0.7699 for StripePy, compared to a median F1c of 0.3554 (Fig. 3L); the median Fowlkes-Mallows index (FMr) is 0.7881, whereas its corresponding value was 0.3765 in the classification (Fig. 3M). While StripePy drastically improves its performance, Chromosight and StripeCaller merely carry over their TPR, PPV, F1c, and FMc values to AHR, FGC, F1r, and FMr. This is because these tools do not have any notion of widths, and thus their metrics are not impacted by interpreting the problem as recognition rather than classification (Fig. 3J-M).

To summarize, estimating the width of a stripe not only provides a more realistic description of the stripe properties, but also significantly improves the performance of a stripe caller, reinforcing the point that stripes should not be thought of as simple 1D segments, but rather as narrow rectangles.

#### StripePy discovers previously detected and novel stripes

We then investigated the degree of overlap between the predictions obtained by StripePy and those produced by Chromosight and StripeCaller.

We compared the anchor sites detected and missed by StripePy with those of Chromosight (Fig. 3N) and StripeCaller (Fig. 3O). The study reveals that, on average, 68.0% (Fig. 3N) to 70.2% (Fig. 3O) of the ground truth anchor points remain undetected by all tools. The vast majority of anchor points correctly identified by Chromosight and StripeCaller, are also found by StripePy: only 4.4% (Fig. 3N) and 2.3% (Fig. 3O) of anchor sites are unique to Chromosight and StripeCaller, respectively. Finally, StripePy recognizes a higher percentage of anchor points, achieving a 18.2% detection rate when StripeCaller is considered (Fig. 3O).

When leveraging stripe widths, StripePy’s advantage increases significantly. The percentage of anchors undetected by StripePy and Chromosight falls to 33.9% (Fig. 3P), for StripePy and StripeCaller drops to 35.5% (Fig. 3Q). Only 2.2% of the anchor points are now unique to Chromosight (Fig. 3P), and just 0.7% are unique to StripeCaller (Fig. 3Q). Meanwhile, a substantial proportion of anchor sites are only found by StripePy: 46.0% compared to Chromosight (Fig. 3P) and 52.9% compared to StripeCaller (Fig. 3Q).

#### StripePy outperforms existing tools across a diverse set of conditions

Hi-C data is generated with varying noise levels and sequencing depths, which affect the resolution at which the data can be meaningfully studied, and significantly impacts data analysis, such as the identification of architectural stripes [35]. Unlike other functional genomics assays, Hi-C data requires analysis at a user-determined effective resolution, often involving the consideration of multiple bin-sizes on the basis of the pattern under study. However, higher resolution increases technical noise and contributes to data sparseness. These factors are influenced, among other things, by the technology used, such the digestion strategy, e.g., sequence-specific digestion by one or more restriction enzymes, or non-sequence specific digestion by MNase (Micro-C) or DNAse (DNAse Hi-C) [33, 40, 54, 64]. To study the effects of contact density, noise, and resolution on the callers, we stratified the dataset by these factors and evaluated the dependency of classification and recognition metrics.

To this end, we grouped contact maps by specifying one factor at a time. For the sake of clarity, here we limit the analysis to the medians of the classification and recognition scores; boxplots of the scores can be found in the Supplementary Data – Supplementary Figs. 2, 3, 6.

Upon first investigation, we observe that medians yield rankings in line with the overall analysis: StripePy consistently outperforms other tools across all contact densities, noise levels, and resolutions for the majority of measures. Focusing on base measures, StripePy provides the highest TPR scores (Supplementary Fig. 1, panels A1, B1, and C1), AHR scores (Supplementary Fig. 1, panels A9, B9, and C9) and FGC scores (Supplementary Fig. 1, panels A10, B10, and C10); however, StripePy exhibits a generally lower median performance in correctly labeling negative bins, as displayed by TNR (Supplementary Fig. 1, panels A2, B2, and C2).

When it comes to PPV, StripePy achieves median values similar to those of Chromosight and StripeCaller at higher contact density and noise level (Supplementary Fig. 1, panels A3, B3). StripePy’s leads in classification and recognition is confirmed when adopting single-value classification (bACC, GM, F1c, FMc, and JI, see Supplementary Fig. 1, panels A4-A8, B4-B8, and C4-C8) and recognition measures (F1r, and FMr, see Supplementary Fig. 1, panels A11-A12, B11-B12, and C11-C12).

We observe that increasing contact frequency does not always result in higher scores. For example, the medians of TPR (Supplementary Fig. 1A1) and AHR (Supplementary Fig. 1A9) show that increasing the contact density results in a slight decrease in StripePy performance. Conversely, Chromosight experiences a minor increase in its performance, while it is only StripeCaller that progressively increases its medians, although it appears to asymptotically approach a value well below StripePy and Chromosight.

Regarding changes in noise level, we see that the median of TPR (Supplementary Fig. 1B1), AHR (Supplementary Fig. 1B9), and of all single-value metrics (Supplementary Fig. 1, panels B4-B8 and B11-B12) show drops of different magnitudes as noise increases. This trend is accompanied by a rise in the median TNR scores (Supplementary Fig. 1B2). The simultaneous rise in median TNR scores and decrease in TPR scores suggests that all callers tend to reduce the number of detected stripes as noise increases, as confirmed by the data in Supplementary Table 1.

With the sole exception of TNR (Supplementary Fig. 1C2) and AHR (Supplementary Fig. 1C9), higher resolutions lead to a drop in median scores for StripePy and Chromosight; the same applies to StripeCaller on a smaller number of metrics. For example, all callers show decreases in median FMc (Supplementary Fig. 1C7) and FMr (Supplementary Fig. 1C12) when transitioning from 10 kb to 5 kb. This might be due, among other reasons, to the increase in sparsity that accompanies the increase in resolution.

#### Analyzing statistical significance

We conducted statistical significance tests to assess whether the variations in classification and recognition performance between pairs of tools are statistically substantial. Supplementary Table 6 presents the *p*-values for the Anderson-Darling test [3], utilizing the implementation provided by SciPy. The tests were conducted on subsets of the StripeBench benchmark, which were obtained by grouping contact maps by specifying one factor at a time, as well as on the entire benchmark. The *p*-values are clamped to a minimum value of 10^−4^. This analysis reveals that, with the sole exception of PPV, the differences between tools are generally statistically significant: in fact, the overwhelming majority of *p*-values fall below the 0.05 threshold, allowing us to reject the null hypothesis that pairs of tools behave according to the same probability distribution. Notable exceptions occur in the classification measures for StripePy and Chromosight, e.g., for low noise level.

### Analyzing human Hi-C matrices with StripePy reveals fast and robust detection of stripes

To investigate how StripePy behaves on real data, we gathered files from the 4DNucleome and ENCODE, and compared the stripes predicted by various available callers at a resolution of 10 kilo-base pair (kbp) with the CTCF peaks from the corresponding cell lines.

We selected four contact maps: an in-situ Hi-C and a Micro-C map from the H1 human embryonic stem cell line (H1-hESC), and an in-situ Hi-C and an intact Hi-C map from the human lymphoblastoid GM12878 cell line. This choice allows us to compare the tools across different cell lines with distinct contact patterns (e.g., sparsity and number of interactions) and technologies, thus providing a comprehensive evaluation of performance under varied conditions. Each interaction map was paired with a CTCF ChIP-Seq dataset. These datasets were used to generate a list of stripe anchor points by taking the midpoint between the start and end coordinates of a CTCF peak. At a resolution of 10 kb, the Hi-C and Micro-C maps for H1-hESC consist of 215,229,365 and 276,917,669 non-zero entries, which sum up to a total of 2,785,926,565 and 1,186,984,847 interactions, respectively (numbers refer to the cis portion of the contact map). This highlights that although the Hi-C map contains more than twice as many interactions when compared to the Micro-C map, it is sparser. As per the intact Hi-C and in-situ Hi-C maps for GM12878, the first has 278,180,961 non-zero entries and a total of 1,415,005,237 interactions, while the second has 91,009,553 non-zero entries and 338,111,285 interactions (numbers refer to the cis portion of the contact map at 10 kb). This indicates that the first map is less sparse and has a higher number of interactions than the second one.

#### Running StripePy on real data confirms its performance in stripe recognition

Since these datasets are also manageable by Stripenn, we have included this tool in the comparative analysis.

Snapshots of neighborhoods from different chromosomes are provided in Supplementary Fig. 5. Supplementary Table 7 reports the classification and recognition measures for the three maps, complete of the number of anchors found (nAF) and the number of stripes predicted (nSP). To ease the comparison, classification and recognition scores are reported as percentages. From plots and statistics we reach a series of conclusions.

First, Chromosight, StripeCaller, and Stripenn focus on visually prominent segments, while StripePy can also locate milder or hidden patterns associated with ChIP-Seq peaks. An example of this is indicated by the black arrows in Supplementary Fig. 5A1-A3, which shows interactions and stripes for the H1-hESC Micro-C dataset. This point is further highlighted by the number of anchors found (nAF), which is significantly higher for StripePy: for the same contact map, stripes predicted by StripePy contain a total of 32,644 anchor sites, compared to 6,257, 5,294, and 9,263 anchors identified by Chromosight, StripeCaller, and Stripenn, respectively (Supplementary Table 7). The higher number of anchors found is accompanied by a significantly higher number of predictions (nSP): for this contact map, StripePy 26,210 candidates, compared to 11,765, 12,018, and 3,864 predictions from Chromosight, StripeCaller, and Stripenn, respectively (Supplementary Table 7).

Moving to the analysis of the classification and recognition measures, we observe that most of the general findings uncovered with the synthetic data from StripeBench are still valid on these maps (Supplementary Table 7). For classification measures, we indeed observe that StripePy is stronger in correctly predicting positives rather than negatives: TPR values are higher in StripePy, while the TNR is generally lower. As for StripeBench, the overwhelming number of non-anchor bins results in an imbalanced classification problem, causing the other methods to focus on the most populous class (i.e., the negatives) at the expense of the anchor sites (i.e., the positive class). When focusing on single-value metrics, StripePy generally achieves the highest scores, making it the best overall classifier. The sole exceptions involve the H1-hESC Micro-C (for bACC and FMc) and GM12878 DpnII in-situ Hi-C datasets (for the sole bACC). A possible explanation for this underperformance is the sparsity of these two maps, which can affect the pseudo-distributions by generating spurious local maxima with little, if any, biological significance. In the most severe case, namely the GM12878 in-situ Hi-C dataset, the sparsity is compounded by a low number of interactions, leading to a significant increase in the number of candidate stripes (see Supplementary Fig. 5D1-D3). For base recognition measures, StripePy excels in AHR but lags behind Stripenn with respect to FGC. The latter can identify however only between 9.10% (in GM12878 in-situ Hi-C) and 17.00% (in H1-hESC Micro-C) of the anchor sites, which contributes to its relatively low overall performance in recognition measures like F1r and FMr. In other words, Stripenn excels in finding a very limited number of stripes which, in turn, are highly likely to contain anchor sites (see Supplementary Fig. 5A4, B4, C4, and D4). Conversely, StripePy has a lower percentage of stripes that contain anchor sites but provides a much more balanced performance in terms of stripe recognition (see Supplementary Fig. 5A1, B1, C1, and D1).

### StripePy exhibits excellent computational performance

While designing and implementing StripePy, special attention was given to computational performance and efficient usage of computational resources. As a result, StripePy is significantly faster than existing tools, being twice as fast as Chromosight and over 66 times faster than Stripenn. Peak memory usage is also much lower than other tools. Overall, processing the .mcool matrix for ENCFF993FGR [22] – a Hi-C dataset with close to two billion interactions at 10 kbp resolution – using 8 CPU cores takes less than 35 seconds and requires below 650 MB of memory. These results were obtained thanks to deliberate decisions while designing the StripePy algorithm, as well as careful implementation taking advantage of various optimization techniques, including parallel programming, asynchronous programming, and exploiting shared memory as much as possible. Furthermore, special care was taken when selecting third-party dependencies. For example, by using hictkpy instead of other Python libraries to read interactions to fetch interactions from .hic and .mcool files, we were able to take advantage of the library capabilities to efficiently fetch interactions surrounding the matrix diagonal, greatly reducing peak memory usage while avoiding needlessly fetching interactions not required by StripePy [57]. On the algorithmic side, unlike other stripe recognition tools, StripePy does not rely on steps rooted in image processing and analysis. While the use of the global pseudo-distribution and the subsequent identification of vertical linear patterns as maximum points resemble the Hough transform [28, 42] – a popular pattern recognition technique for curve and surface recognition – avoiding the use of binarization and edge detection allows to speed up the computation and eliminates the need for additional parameters.

## Discussion

Hi-C and related techniques have greatly advanced our understanding of 3D genome organization and its role in gene regulation, DNA replication, and repair [26, 59]. While computational methods have been crucial to these insights [60], tools for detecting architectural stripes remain scarce. Here, we developed StripePy, a new CLI application which combines concepts from geometric pattern recognition, algebraic topology, and basic geometry to detect architectural stripes from Hi-C data. StripePy can process contact maps in various formats (.cool, .mcool, and .hic) and outputs its findings in both Hierarchical Data Format (.hdf5) and BEDPE format. Unlike most existing solutions, StripePy does not reduce stripe detection to a mere bin classification problem. Instead, it also computes shape descriptors such as the width and height of stripes, and generates statistics that can be used for postprocessing purposes, such as ranking and classification of candidate stripes to, e.g., filter out unrealistic stripes.

A major challenge in the field is the lack of standardized definitions for architectural elements in Hi-C data, such as TADs, stripes, and individual contacts [8, 63, 37, 52]. To address this, we developed StripeBench, a dataset of 64 simulated contact maps with ground truth annotations and a comprehensive list of metrics, facilitating exhaustive quantitative comparisons between callers. We anticipate that this benchmark will aid in the future development of algorithms tailored for Hi-C and related data.

An additional obstacle with the analysis of Hi-C data lies in the variation in resolutions, noise levels, and sequencing depth that characterize this kind of data, all of which hinder standardized detection of chromatin features. Our benchmark tackles this issue by incorporating datasets with diverse conditions, enabling a more comprehensive evaluation of stripe detection methods.

Based on the benchmarks presented in the previous sections, it is clear from the quantitative analysis that all tools struggle, to different degrees, in accurately identifying true positives in the classification task. This difficulty is mainly caused by the skewed distribution of the ground truth labels, but it also stems from the lack of a precise and unambiguous definition of what a stripe is. While incorporating geometric descriptors proves beneficial – as noticeable from the increased F1-scores and Fowlkes–Mallows indexes – further work is needed to better define architectural stripes and their functional implications.

## Acknowledgments

We thank the 4DNucleome Network and the lab of Job Dekker for contributing several of the Hi-C and Micro-C datasets used as part of this study.

We thank the ENCODE Consortium and the labs of Erez Aiden and Michael Snyder for contributing with ENCODE data used as part of this study.

Furthermore, we would like to acknowledge Mr. Bendik Berg for his contributions in packaging StripePy and configuring the framework for unit testing.

## Funding

This work was supported by the Norwegian Research Council projects 324137 and 343102.

## Code availability

StripePy source code is hosted on GitHub at https://github.com/paulsengroup/StripePy and is archived on Zenodo at DOI: 10.5281/zenodo.15310827 [58]. StripePy can be easily installed from source or PyPI using pip https://pypi.org/project/stripepy-hic. Furthermore, StripePy is available on Bioconda at https://anaconda.org/bioconda/stripepy-hic and can be installed using conda. Containerized versions of StripePy are regularly published on DockerHub at https://hub.docker.com/r/paulsengroup/stripepy.

Code used for the benchmarks and data analyses presented in this article are hosted on a separate GitHub repository at https://github.com/paulsengroup/2024-stripepy-paper. A copy of the code in this repository is archived on Zenodo at DOI: 10.5281/zenodo.15310693 [53].

## Data availability

This study re-analyzed a number of publicly available datasets released by the 4DNucleome [13, 55] and ENCODE [17, 29, 38]. Hi-C and Micro-C datasets: 4DNFI6HDY7WZ [33, 44] - H1-hESC (in situ Hi-C; DpnII); 4DNFI9GMP2J8 [33, 45] - H1-hESC (Micro-C); ENCFF216QQM [19] - GM12878 (in situ Hi-C; DpnII); ENCFF993FGR [22] - GM12878 (intact MNase Hi-C).

CTCF ChIP-Seq datasets: ENCFF692RPA [20] - H1-hESC; ENCFF796WRU [21] - GM12878.

The results presented throughout the manuscript have been deposited on Zenodo at DOI:10.5281/zenodo.15308825 [48].

The benchmark developed as part of this publication and used to contrast the stripe callers considered in this manuscript, StripeBench, has also been deposited on Zenodo at DOI:10. 5281/zenodo.14448329 [47].

**Andrea Raffo**, PhD, is a postdoctoral candidate in Bioinformatics at the University of Oslo, having obtained his PhD in Mathematics from the same institution in 2022. His research interests revolve around geometric modeling and shape analysis, with focus on various application domains such as biosciences, computer graphics, and Computer-Aided Design.

**Roberto Rossini**, PhD, is a postdoctoral candidate in Bioinformatics at the University of Oslo. His research efforts focus on the field of 3D-Genomics, where he develops and applies novel computational tools and analysis workflows.

**Jonas Paulsen**, PhD, is Professor at the Department of Biosciences at the University of Oslo. With a significant cross-disciplinary focus spanning computational and life sciences, his research involves the development of computational methods and analyses to understand the structure and function of genomes in three dimensions (3D). He is an active member of the International Nucleome Consortium and a contributor to the 4DNucleome Program.

## Footnotes

1 https://github.com/XiaoTaoWang/StripeCaller

2 https://github.com/ChristopherBarrington/domainClassifyR

## Notes

### Competing Interest Statement

The authors have declared no competing interest.

### Summary of Updates

We conducted an Anderson-Darling test to assess the statistical significance of differences in classification and recognition performance, both across the entire StripeBench benchmark and when grouped by individual factors; a new section has been added to discuss computational performance.

