## Supplementary Information for "StripePy: fast and robust characterization of architectural stripes"

#### Extended description of StripePy

StripePy runs through four consecutive steps: a pre-processing (Step 1), followed by the identification of genomic bins exhibiting linear patterns (Step 2), the subsequent estimation of geometric descriptors (Step 3), and the final definition of biological descriptors (Step 4). A compact overview of the algorithm is given in Fig. 1, while a more detailed representation is provided in Supplementary Fig. 7.

##### Step 1: Laying the groundwork for data exploration

To mitigate the bias of the *genomic distance effect* – namely the tendency of higher prevalence of crosslinks between genomic loci close together along the genome even in the absence of any specific higher order structure – we perform global enhancement via the well-known log-transformation. The log-transformation is a global intensity transformation which expands narrow ranges of low intensity values and compresses wide ranges of high intensity values. The general form of the log-transformation is

$$P(x, y) = c \log(1 + H(x, y)), \tag{1}$$

being $H(x, y) \geq 0$ the initial intensity at bin pair $(x, y)$, $P(x, y)$ the output intensity at the same location, and $c$ a constant. Here, we set $c$ so that intensities are projected onto $[0, 1]$. Furthermore, we retain only entries that fall within a specific band around the main diagonal (default bandwidth: 5 Mbp), as we are specifically targeting patterns anchored at the main diagonal. As an example, Supplementary Fig. 7A-B provides a graphical illustration of how global enhancement acts on a specific square sub-region along the main diagonal (what we will call a *region of interest*).

We proceed by splitting the pre-processed Hi-C matrix $P$ into lower- and upper-triangular matrices, henceforth denoted as $L$ and $U$, respectively; see Supplementary Fig. 7C-D.

#### Step 2: Line detection via pattern recognition and Topological Data Analysis

For the sake of conciseness, we here adopt two simplifications. First and foremost, we restrict our explanation on the lower-triangular matrix, i.e., $L$ – as the processing of the upper-triangular part $U$ is analogous; for visualization purposes, however, we will often show the full Hi-C matrix $P$. We choose to display the full matrix as it can help the visual analysis of an algorithm's output, as any vertical pattern can be found horizontally given that Hi-C matrices are naturally symmetric. Secondly, we limit ourselves to recognizing vertical patterns, again because of symmetry.

We sum $L$ over the rows, thus obtaining a scalar function, or 1D profile, which can be interpreted as a *pseudo-distribution* $\pi_{\text{glob}}$. The function is smoothed by using weighted quasi-interpolation [7–9], a weighted moving average technique for curve approximation. A graphical example is given in Supplementary Fig. 7E. The term "pseudo" used in this context is necessary as $\pi_{\text{glob}}$ is not, strictly speaking, a probability distribution. The location of each local maximum points at a possible linear pattern. Note that the pseudo-distribution bears resemblance to the accumulator function derived by applying the Hough Transform [1, 4], but in this case the search for vertical linear patterns is conducted without any edge detection or binarization to speed up computation and avoid the introduction of additional parameters.

We filter local maxima by measuring their significance via *topological persistence*. In the analogy of the water level, the pseudo-distribution is a submerged bank. Imagine the water level to gradually go down: when it reaches a local maximum, a new island is formed, and its height is called *birth level*; when it reaches a local minimum, two islands merge. The higher island is, by convention, considered to subsume the lower island, and the local minimum is the lower island's *death level*. The topological persistence of an island, i.e., its significance, is the difference between its birth and death levels. The local maximum points of $\pi_{\text{glob}}$ that survive the filtration process will be here referred to as the *seed sites* of candidate stripes, and will be denoted as $s_1 < s_2 < \cdots < s_n$. An example of seed sites is provided in Supplementary Fig. 7F, while the corresponding locations are superimposed to the contact map in Supplementary Fig. 7G.

To avoid spurious seed sites from sparse regions, we apply the following additional check. For each seed site, we consider a neighborhood of pre-defined size and calculate the ratio of bins – within this neighborhood – where $\pi_{\text{glob}}$ exceeds 0.1 (i.e., 10% of the maximum height); if this ratio is greater than 85%, the seed site is retained, otherwise it is discarded.

#### Step 3: Shape analysis

Once the seed sites (i.e., sites hosting linear patterns) $s_1, \ldots, s_n$ are detected in the lower-triangular preprocessed Hi-C matrix $L$, we proceed by estimating width and height of each candidate stripe $S_j$ independently as follows.

**Width estimation** We start by defining a sub-interval $I_j := [i_j^L, i_j^R] \subset I$ so that $I_j$ is the largest interval with the following properties: (1) $s_j$ is contained in $I_j$; (2) $\pi_{\text{glob}}$ is monotonically increasing in $[i_j^L, s_j]$; (3) $\pi_{\text{glob}}$ is monotonically decreasing in $[s_j, i_j^R]$. To define the width of of $S_j$, we look at the global pseudo-distribution $\pi_{\text{glob}}$ in the interval $I_j$. The right boundary of the candidate stripe is defined as

$$b^R(s_j) := \min\{\tilde{b}^R(s_j), b + w_{\max}/2\}, \tag{2}$$

where

$$\tilde{b}^R(s_j) := \arg\max_b \left\{ \pi_{\text{glob}}(b) - \pi_{\text{glob}}(b-1) \middle| s_j < b \leq i_j^R \right\}; \tag{3}$$

the left boundary is given by

$$b^L(s_j) := \max\{\tilde{b}^L(s_j), b - w_{\max}/2\}, \tag{4}$$

where

$$\tilde{b}^L(s_j) := \arg\max_b \left\{ \pi_{\text{glob}}(b+1) - \pi_{\text{glob}}(b) \middle| i_j^L \leq b < s_j \right\}. \tag{5}$$

Here, $w_{\max}$ denotes the maximum width, which by default is set to a value of 100 kb.

These formulas correspond to defining the boundaries as the genomic bins exhibiting largest values of (approximate) first-order derivative of $\pi_{\text{glob}}$ in a left or right neighbourhood of $s_j$, with a maximum width as additional constraint. An exemplification of this description is shown in Supplementary Fig. 7H. Given a seed site, the left and right boundaries naturally define horizontal intervals, sometimes referred to as horizontal domains: see Supplementary Fig. 7I for a graphical display of what the single horizontal domains mean in terms of (lower-triangular) matrix slicing.

The width of a candidate stripe is computed by $b^R(s_j) - b^L(s_j)$.

**Height estimation** Given the $j$-th seed site and the corresponding horizontal domain made of left and right boundaries, we define a vertical domain (lower and upper boundaries) by introducing a *local* pseudo-distribution. Such a profile is obtained by applying the following procedure (also see Supplementary Fig. 7I-J):

- Slice $L$: filter out columns that do not belong to the current horizontal domain; discard rows from the upper-triangular part of the matrix.
- Sum (i.e., marginalize) over the surviving columns, then scale by dividing by the global maximum. Smooth the local pseudo-distribution by using, again, the weighted quasi-interpolation.

Two criteria are available: (1) By applying topological persistence to find persistent maxima, we can determine where the local 1D pseudo-distribution presents significant peaks; we use the peak that is farthest from the main diagonal to define the stripe boundary. (2) The vertical domain of a candidate stripe is retrieved by finding the smallest interval containing all coordinates where the smooth approximation is above a user-threshold. When studying a candidate stripe in the lower-triangular matrix, the top-boundary $b^U(s_j)$ is naturally anchored to the main diagonal; for bottom-boundary $b^D(s_j)$, the steepest the local pseudo-distribution, the closer $b^D(s_j)$ is to the main diagonal. The height of a stripe is then computed as $b^D(s_j) - b^U(s_j)$.

At this point, each quintuplet consisting of seed site, left-, right-, top-, and bottom-boundaries, defines uniquely a candidate stripe from a geometric viewpoint. We will denote candidate stripes as $S_1, \ldots, S_n$ to distinguish them from seed sites $s_1, \ldots, s_n$; an example is shown in Supplementary Fig. 7K.

#### Step 4: Signal analysis

To enrich a purely geometric structure with biological information, we compute descriptors that summarize information about the signal distribution – both inside and outside of candidate stripes. To prevent our descriptors from being influenced by the genomic distance effect, we apply a correction to the shape of a stripe. Instead of selecting a number of rows equal to the stripe's height, we select a number of diagonals matching its height, resulting in a parallelogram, see Supplementary Fig. 7L.

- *Inner descriptors* We adopt the five-number summary, a set of descriptive statistics consisting of five sample percentiles:
  - The minimum, sometimes called $Q_0$ or 0th percentile.
  - First ($Q_1$), second ($Q_2$), and third ($Q_3$) quartiles – which consists of the 25th, 50th, and 75th percentile, respectively. The second quartile corresponds to the median; the first and third quartiles correspond to the median of the lower and the upper half of the dataset, respectively.

- The maximum, sometimes called $Q_4$ or 100th percentile.

In addition, we consider the arithmetic mean and standard deviation of the signal within the stripe boundaries. We will call the latter as *mean inner signal*.

- *Outer descriptors.* We compute the statistics from the previous point using regions that neighbor the candidate stripe. More precisely, the *left k-neighbour* $\mathcal{N}_j^R$ consists of the first $k$ diagonals to the left of $S_j$; analogously, the *right k-neighbour* $\mathcal{N}_j^R$ consists of the first $k$ diagonals to the right of $S_j$. We will denote as $\mathcal{N}_j := \mathcal{N}_j^L \cup \mathcal{N}_j^R$ the *k-neighbour* of the $j$-th candidate stripe. The arithmetic mean of the signal inside the $k$-outer neighbour will be referred to as *mean k-outer signal.*

- *Mixed descriptors.* We define the *k-relative change* as the difference between mean inner and mean $k$-outer signals, divided by the mean $k$-outer signal. The relative change resembles the estimation of a first-order derivative at the stripe boundary, when performed by using a first-order finite difference method.

### Extended description of StripeBench

This section is devoted to provide additional information on the StripeBench benchmark, intended as a dataset equipped of a ground truth and a set of performance measures.

#### Dataset

To generate Hi-C matrices we consider a genome-wide run of MoDLE with loop extrusion simulated by using barriers imputed from RAD21 ChIP-Seq H1-hESC data, and with default settings except for the following three factors:

- Target contact density $\delta$, i.e., the average contact density to be reached before the simulation is halted, here set to 1, 5, 10, and 15.

- Scale parameter $\sigma$ of the generalized extreme value distribution, used to randomize contacts by additively introducing noise to molecular contacts, is set to 0, 5000, 10000, and 15000.

- Resolution $r$ in base-pairs for the output contact matrix in cooler format, set to 5k, 10k, 25k and 50k.

The values chosen for the three factors lead to 16 Hi-C matrices, each one available at four resolutions for a total of 64 simulated matrices. The choice of target contact density and scale parameters in our study is driven by the objective of considering Hi-C matrices with diverse sequencing depths and increasing levels of noise, while still maintaining a manageable number of matrices. For a graphical illustration of the effects of these parameters on the contact maps, the reader is referred to Fig. 2A.

#### Ground truth

For each Hi-C matrix, the ground truth is obtained by retrieving loci with average extrusion barrier occupancy greater or equal than 0.70. Extrusion barrier occupancy consists of a scalar in $[0, 1]$, and is therefore interpretable as a probability value; intuitively, the higher this value, the more visible the stripe is, as shown in visual examples in Fig. 2B.

We then proceed by merging extrusion barriers mapping to the same bin at a given resolution – so that each locus appears only once; however, we consider the lower- and the upper-triangular parts of the contact map separately, so that a bin can yet be deemed relevant for both parts of the matrix, either individual parts or neither of them. Each of loci will be henceforth referred to as *anchor site* or *anchor point*.

#### Different measures for different problems

The capability of a tool to find meaningful stripes is here quantified by using classification and recognition measures. We hereby adopt the following terminology:

- *Classification* is the task of assigning labels to input data, such as categorizing genomic bins as either 0 or 1 based on whether or not they contain an anchor point. We therefore discard the notion of width and evaluate a method's ability to identify whether a certain bin *hosts (i.e., contains)* an anchor point or not. For Chromosight and StripeCaller, obtaining the predicted classification vector is straightforward since these methods specifically set the stripe width to the resolution of the Hi-C matrix[1]; in the case of StripePy, we obtain the predicted classification vector by considering the coordinates of seed points as anchor point predictions; for Stripenn we consider the midpoint of each horizontal interval, since the method does not provide additional information to help picking a location.

- *Recognition* deals with the identification of rectangular regions[2] that host anchor points.

We decide to employ both formulations since neither of them alone is adequate for analysing the problem. Solely relying on recognition is insufficient to assess a set of tools because any partition of the Hi-C matrix can capture all anchor sites with no difficulty, albeit with a potentially excessive number of wide stripes. This limitation originates because it is not possible to obtain a ground truth notion of width or length. On

[1]These two methods return pairs of the form $\{i, i + r\}$, where $r$ is the resolution and $i$ is a genomic coordinate. We consider the bin of genomic coordinate $i$ to host the anchor point.

[2]What stripes are, geometrically speaking

the other hand, classification measures does not account that stripes are not unidimensional segments, but rather narrow rectangles.

The following "base" measures are adopted for the classification problem:

- *Sensitivity and Specificity* measure the ability of a method to correctly identify positives and negative cases, respectively. By *positives* we here mean bins that are flagged as anchor points, while *negatives* are bins that do not host any anchor. Sensitivity is also known as *True Positive Rate* (TPR) or *recall*, while specificity goes under the name of *True Negative Rate* (TNR).

- *Positive Predictive Value* (PPV), or *precision*, is the proportion of true positives to the total number of items classified as positives by the model.

Similarly, we introduce the following "base" recognition measures:

- *Anchor Hit Rate*, or AHR, computed as the fraction of anchor sites that lie in one of the candidate stripes. It is closely related to TPR, although we are here more permissive since we consider the estimated stripe width in its computation.

- *Fraction of Good Candidates*, or FGC, computed as the fraction predicted stripes hosting an anchor site. It resembles to PPV when accounting for the estimated stripe width.

It is worth noting that there is no metric corresponding to specificity, as we are only able to draw conclusions on candidate stripes (whether these are true or false positives) since we lack a notion of negative.

There are several metrics that are used to handle datasets with a severe skew in the class distribution. The following single-value metrics are adopted:

- *Balanced accuracy* and *Geometric Mean* are single-value metrics that combine sensitivity and specificity by computing their arithmetic or geometric mean. We denote them as bACC, and GM, respectively.

- *F1-score* and *Fowlkes–Mallows index*, here respectively denoted as F1c and FMc, are the harmonic and geometric means of precision and recall.

- *Jaccard Index* (JI), or Jaccard similarity coefficient, measures the similarity between true labels and predicted labels by taking the ratio between the number of true positives against the sum of correctly predicted positive samples and all incorrect predictions. This metric considers both false positives and false negatives in a balanced manner, excluding only the correctly predicted negative samples. It is particularly well-suited for detecting rare events, as it ensures that the evaluation is sensitive to the correct classification of infrequent positive instances without being dominated by numerous correct identifications of negative class instances.

In a similar manner, recognition measures are summarized by adapting F1-score and Fowlkes–Mallows index: in this adaptation, TPR is replaced by AHR and PPV is replaced by FGC. We denote them as F1r and FMr, respectively.

### Benchmarking analysis

The following tools were installed for the comparative analyses: Chromosight version 1.6.3 [3], StripeCaller version 0.1.0 https://github.com/XiaoTaoWang/StripeCaller, Stripenn 1.1.65.18 [12], and StripePy version 1.0.0.

### Matrix normalization affects StripePy performance

We decided to investigate the effect of contact map normalization on the performance of StripePy. To this end, maps from Supplementary Table 9 were normalized using hictk [11] with two algorithms: genome-wide iterative correction and eigenvector decomposition (ICE) [2] and genome-wide SCALE [10]. To ensure a consistent comparison, StripePy was rerun on the normalized contact maps with all thresholds unchanged. Snapshots of neighborhoods are provided in Supplementary Fig. 6, while classification and recognition measures are reported in Supplementary Table 9.

We first note that applying normalization significantly reduces the number predicted stripes (nSP). This reduction is visually evident when comparing the plots for the case of no normalization (Supplementary Fig. 6 A1, B1, C1, and D1) with those for ICE- (Supplementary Fig. 6 A2, B2, C2, and D2) and SCALE-normalized maps (Supplementary Fig. 6 A3, B3, C3, and D3). This observation is further confirmed by the nSP values in Supplementary Table 9: for example, for the H1-hESC Micro-C map, nSP decreases from 26,210 to 6,303 and 8,819 for raw, and ICE- and SCALE-normalized maps, respectively. This is not surprising, as one of the effects of Hi-C matrix normalization is smoothing of the original data, resulting in smoothed global pseudo-distributions and, consequently, a smaller number of peaks (i.e., stripe seeds).

Other interesting observations can be made from Supplementary Table 9. First, the decrease in nSP is accompanied by a decrease in the number of found anchors (nAF): for the same map, nAF decreases from 32,644 to 14,722 and 18,708 with ICE- and SCALE-normalized maps, respectively. Moving to the analysis of classification and recognition measures, we observe that for all contact maps, applying normalization increases some base metrics (TNR and FGC) while decreasing others (TPR, PPV and AHR). By examining single-value metrics, it becomes evident that the gain in the former scores is not sufficient to compensate the drop in the latter metrics, resulting in a unanimous decrease in all single-value statistics. Therefore, we conclude that normalizing contact maps before using StripePy for stripe calling might impair performance and should be approached with caution.

#### Impact of the two filtering strategies

StripePy employs two filtering strategies. The first filtration takes place in Step 2 (see Methods), where local maximum points of the global pseudo-distribution are filtered by thresholding their topological persistence values. By applying this filtration to the global pseudo-distribution, we retain genomic bins with a stronger (uni-dimensional) linear pattern, as this process resemble the voting procedure from the Hough transform [1]. The effect of increasing thresholds on these local maximum points is illustrated in Supplementary Fig. 8A-C: in practice, requiring higher topological persistence prioritizes local maxima with greater height differences compared to their surrounding areas. Supplementary Fig. 8 D-E show the candidate stripes produced at the end of Step 3 (see Methods), where each seed site is enriched with estimates of stripe width and height. While acting on topological persistence already reduces the number of candidate stripes, it doesn't account for stripes being two-dimensional patterns. Therefore, a second filtering step is applied, which thresholds the relative change parameter – a score that compares the average signal inside and in the near proximity of a candidate stripe. The effect of a more stringent filtering on this parameter for the set of candidates of Supplementary Fig. 8E is displayed in Supplementary Fig. 8 G-I. As expected, the higher the threshold, the fewer candidates are returned.

Default values for these thresholds are provided with the software to simplify the user's work and enable a fair comparison with alternative stripe callers. However, optimal results may require adjusting these parameters based on the specific contact map. It is often best to run the tool on a chromosome and fine-tune the parameter values on a case-by-case basis to achieve the most accurate results.

#### Pre-processing of interaction matrices released by ENCODE and the 4DNucleome

Matrices in .mcool format published by the 4DNucleome are generated using a version of Cooler that leads to data corruption when generating the .mcool files by iterative coarsening a .cool file [3]. Thus, prior to using dataset 4DNFI6HDY7WZ and 4DNFI9GMP2J8 [5, 6] for any data analysis or benchmark, we restored the corrupted files using `hictk fix-mcool` [11].

[3]Cooler data corruption: https://hictk.readthedocs.io/en/stable/file_validation.html#cooler-index-corruption

Next, interaction matrices in .hic format were converted to .mcool format using `hictk convert` [11].

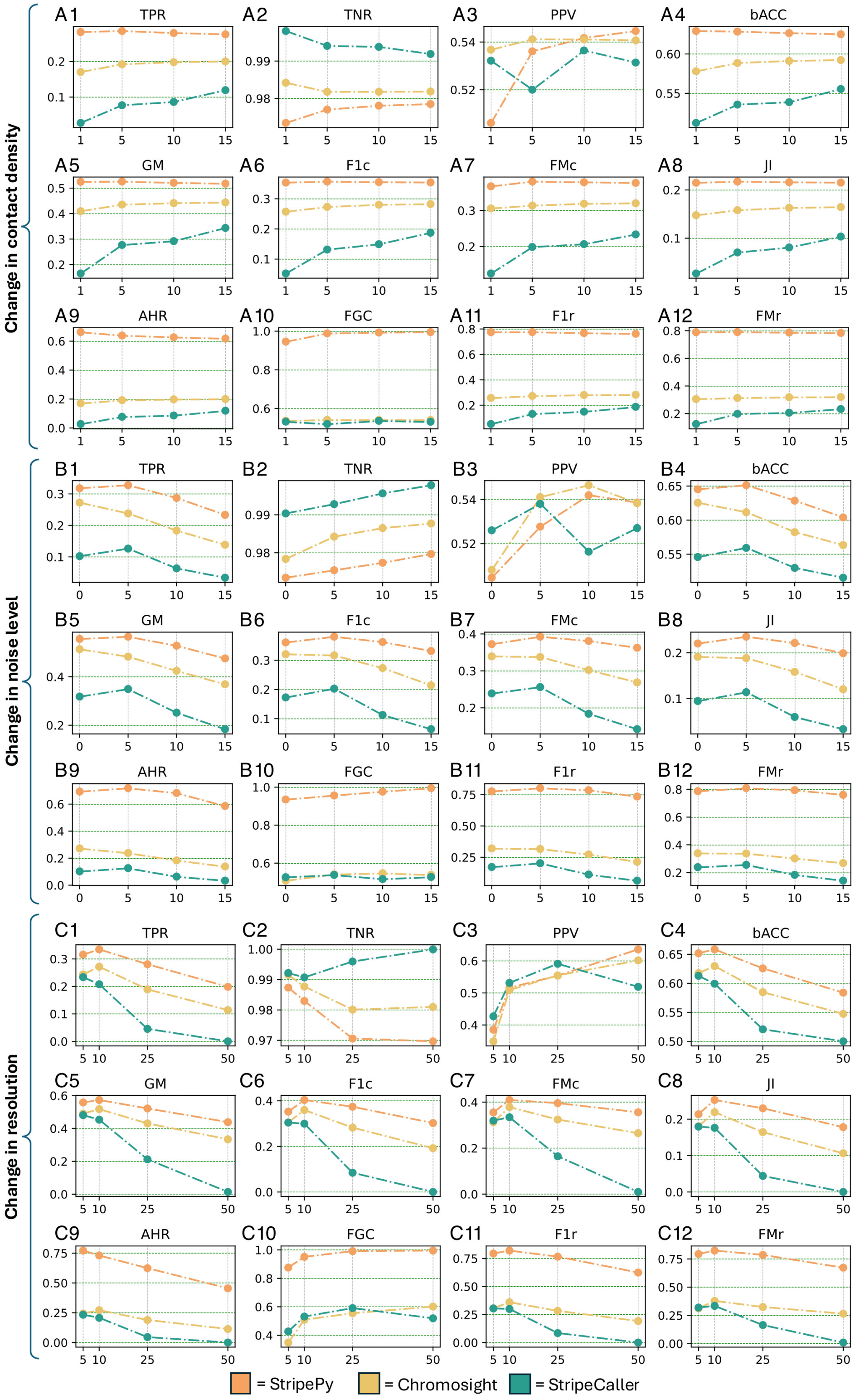


**Supplementary Fig. 1. | Benchmarking pt.2**. Plots of the medians of the classification and recognition measures by single factors: contact density (**A1-A12**), noise level (**B1-B12**), and resolution (**C1-C12**).

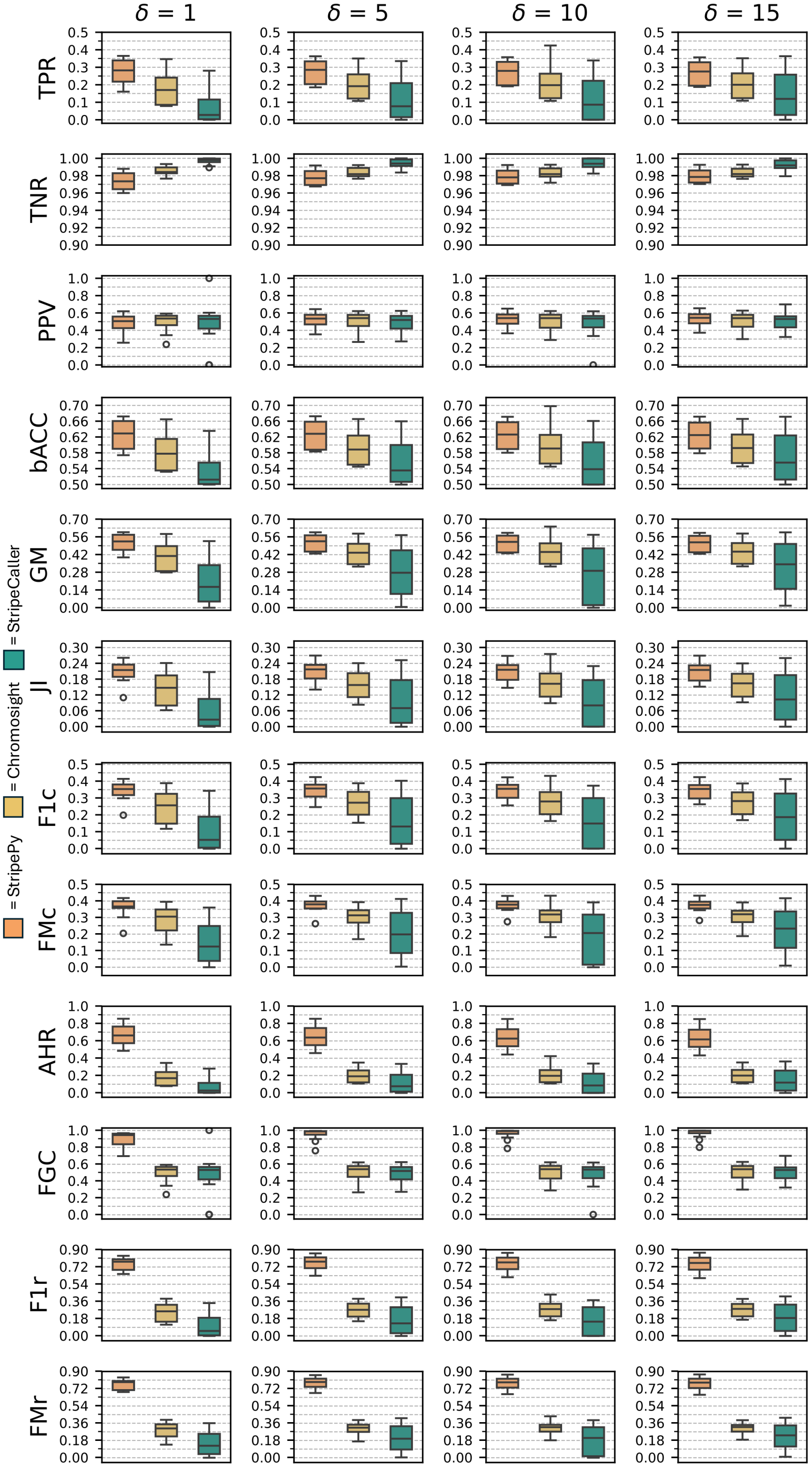


**Supplementary Fig. 2. | Change in contact density for the StripeBench's benchmark**. Boxplots of the classification (TPR, TNR, PPV, bACC, GM, JI, F1c, and FMc) and recognition (AHR, FGC, F1r, and FMr) measures by change in contact density.

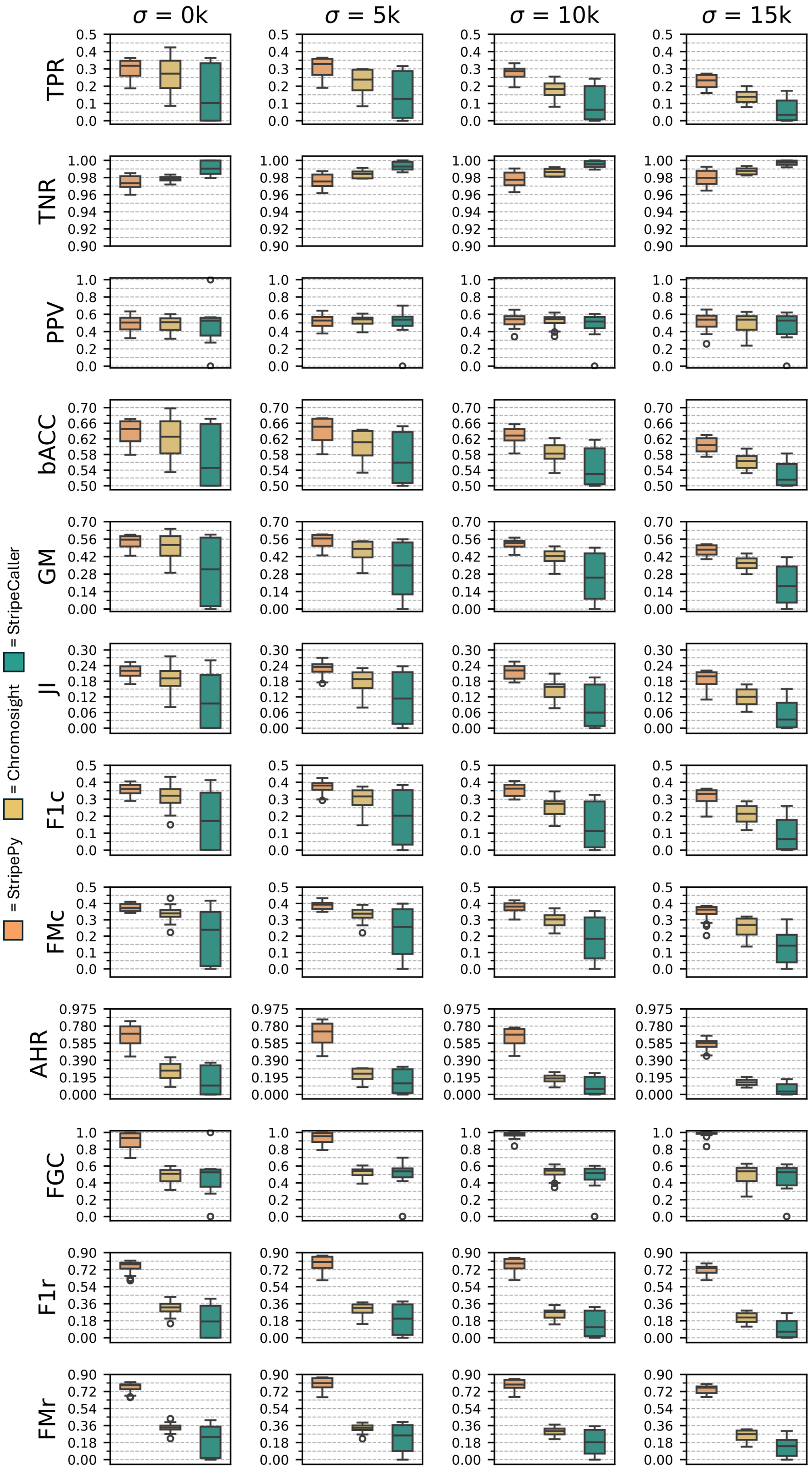


**Supplementary Fig. 3. | Change in noise level for the StripeBench's benchmark**. Boxplots of the classification (TPR, TNR, PPV, bACC, GM, JI, F1c, and FMc) and recognition (AHR, FGC, F1r, and FMr) measures by change in noise level.

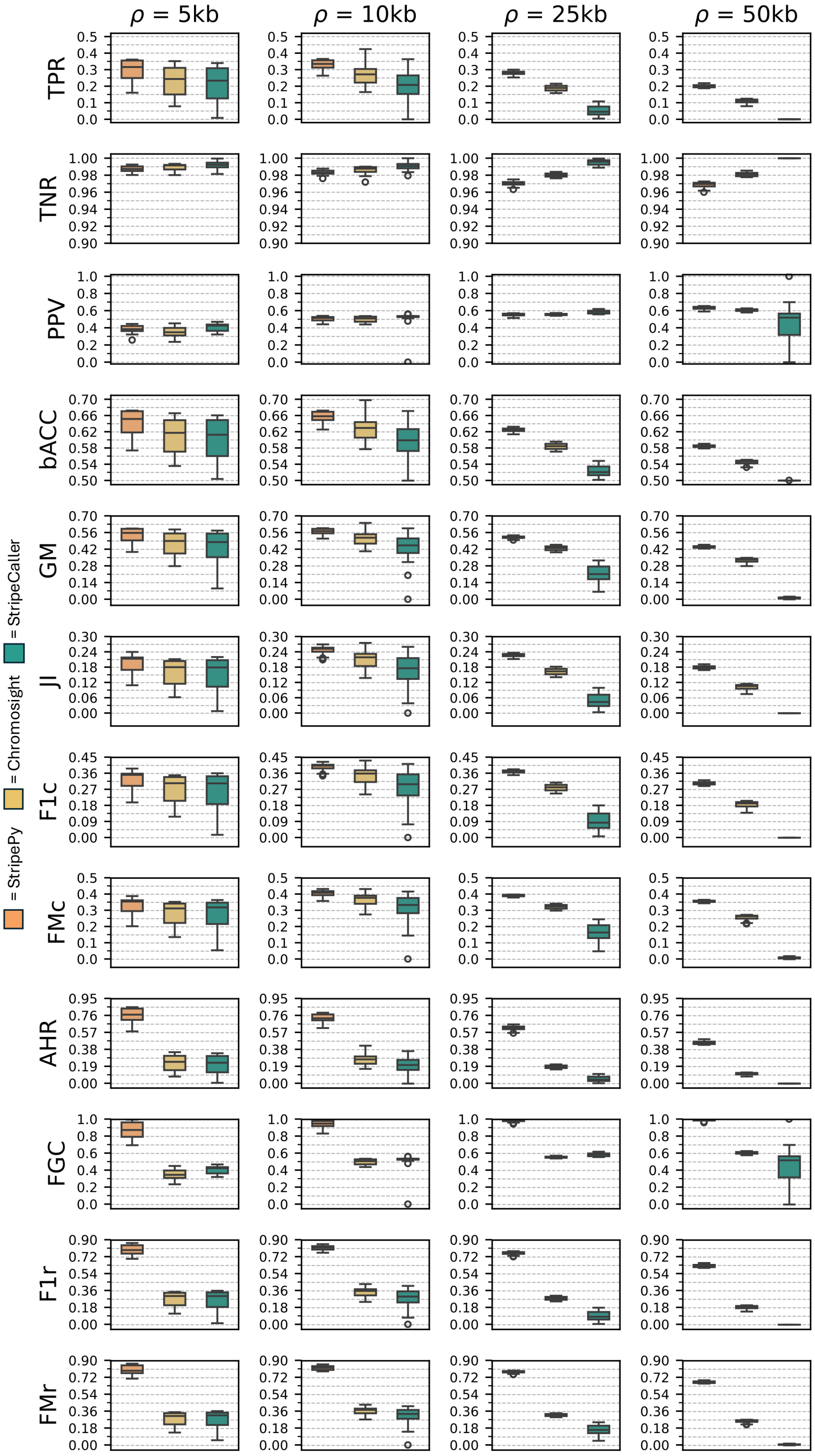


**Supplementary Fig. 4. | Change in resolution for the StripeBench's benchmark**. Boxplots of the classification (TPR, TNR, PPV, bACC, GM, JI, F1c, and FMc) and recognition (AHR, FGC, F1r, and FMr) measures by change in resolution.

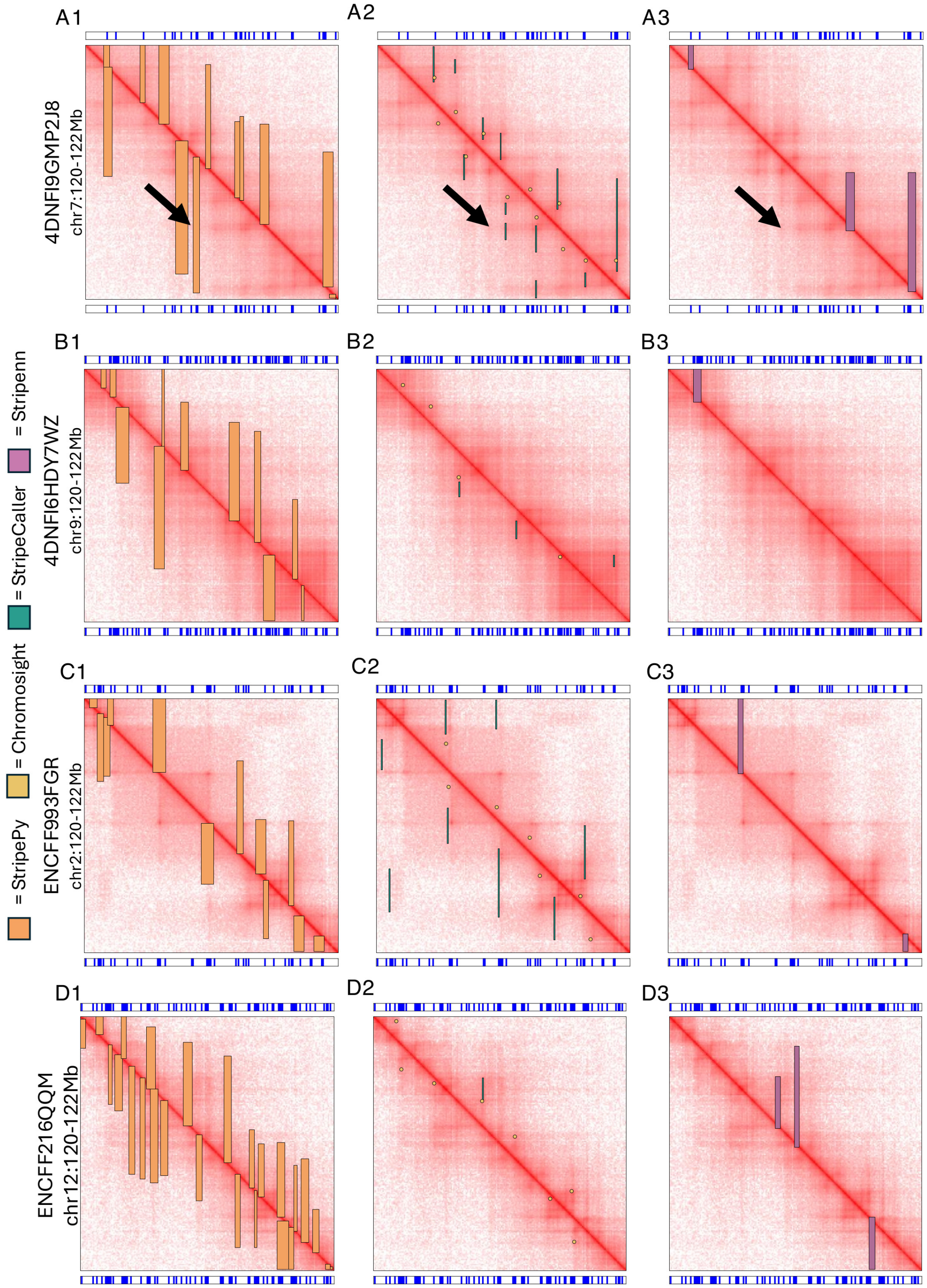


**Supplementary Fig. 5. | Four regions from as many contact maps** . The upper and lower bands point at bin hosting CTCF sites, obtained from ChIP-Seq data. The images of the contact maps have superimposed the predictions produced by the tools under study. For each map, the region is repeated three times to improve clarity: first column consists of StripePy, second column refers to Chromosight and StripeCaller, last column has Stripenn. In the first row, black arrows highlight an example of mild pattern that is correctly identified by our method but overlooked by the other tools.

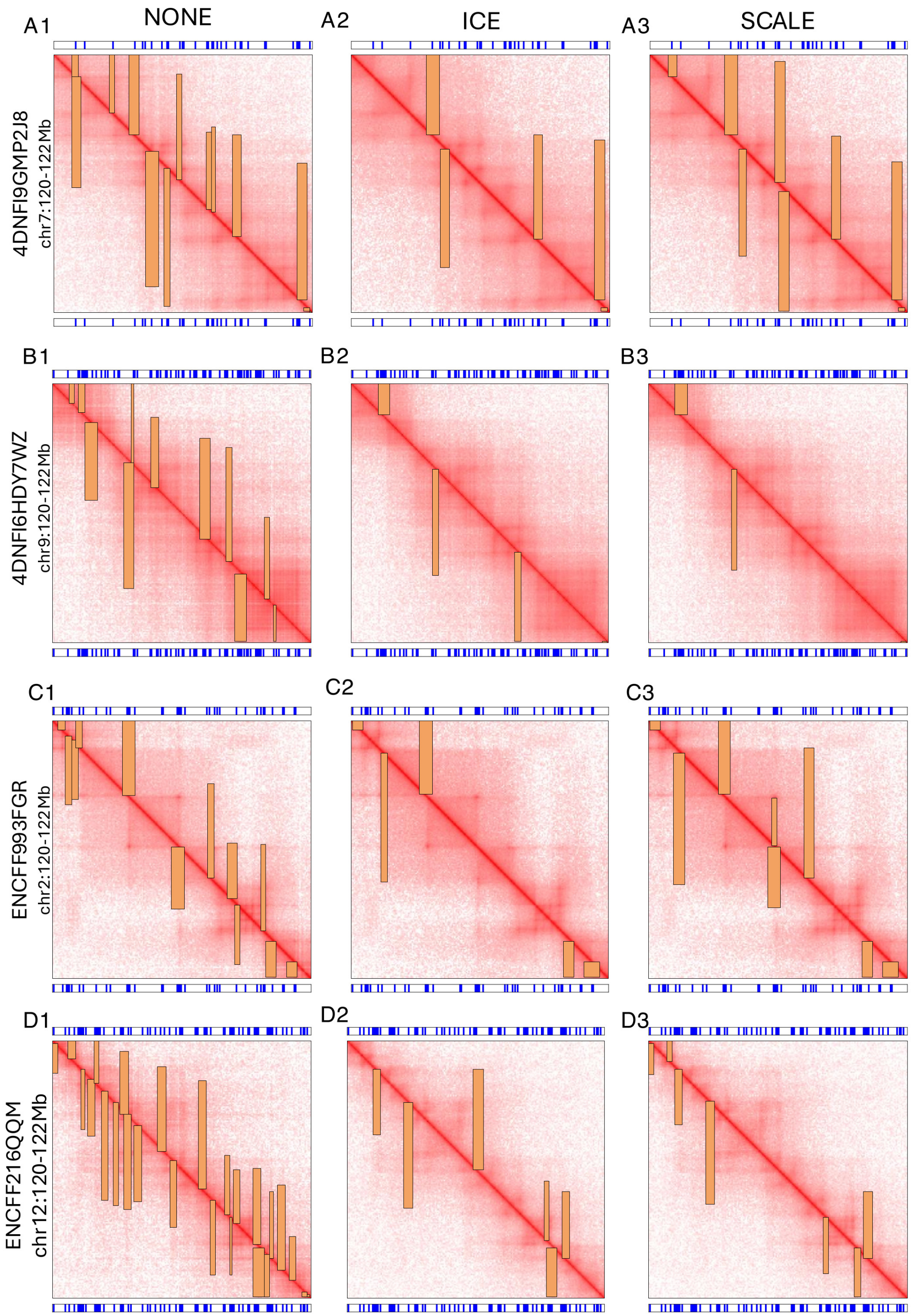


**Supplementary Fig. 6. | Effects of normalization**. The plots represent four regions from as many contact maps. The upper and lower bands point at bin hosting CTCF sites, obtained from ChIP-Seq data. The images of the contact maps have superimposed the predictions produced by StripePy when applied to three conditions: no normalization (NONE), ICE and SCALE. For each map, the region is repeated three times to improve clarity: from left to right, columns refer to NONE, ICE, and SCALE.

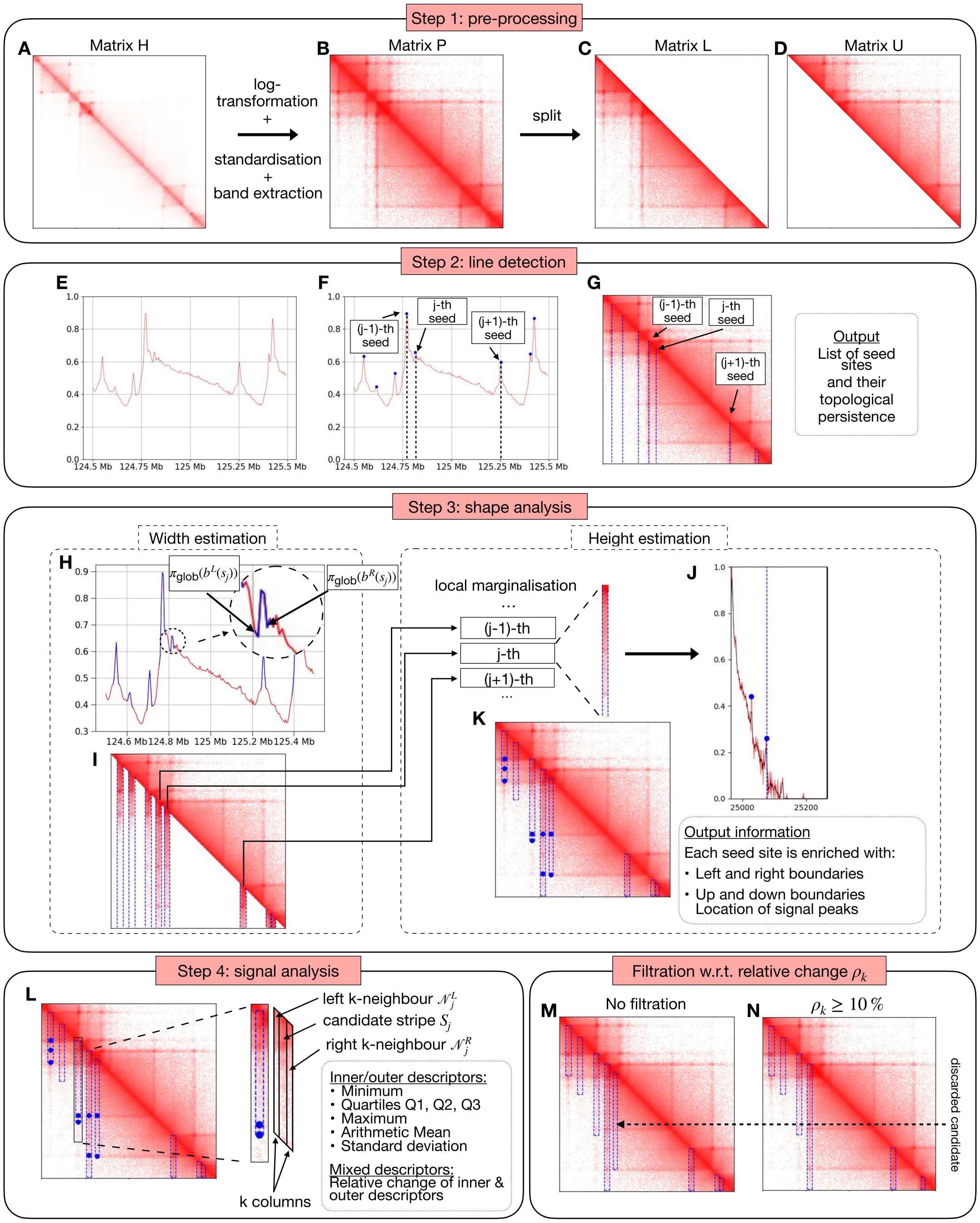


**Supplementary Fig. 7.** | **Extended overview of StripePy**. Step 1: **A** region of an input contact map as it is, **B** after global enhancement via log-transformation, and **C-D** final split into lower- and upper-triangular parts; the remaining steps focused on on the lower-triangular part, to keep the plots readable. Step 2: **E** constrained pseudo-distribution obtained by marginalizing L; **F** constrained pseudo-distribution with superimposed persistent maximum points, a.k.a. seeds; **G** pre-processed Hi-C matrix with superimposed vertical lines corresponding to persistent seeds. Step 3: **H** study of neighbourhoods of seed sites allow to introduce notions of left and right boundaries, which can be then used to slice the pre-processed Hi-C matrix **I**; **J** local pseudo-distributions (original: red curve, smoothed: black curve) allow to introduce up/down boundaries (dashed blue line), as well as to detect the location signal peaks (blue dots); **K** overlap rectangles corresponding to candidate stripes, together with location of signal peaks, to the pre-processed Hi-C matrix. Step 4: **L** shows a region neighbouring a candidate stripe and list the sets of inner/outer/mixed descriptors. The initial set of candidate stripes, represented in **M**, is reduced by thresholding one of these descriptors, namely the relative change parameter **N**.

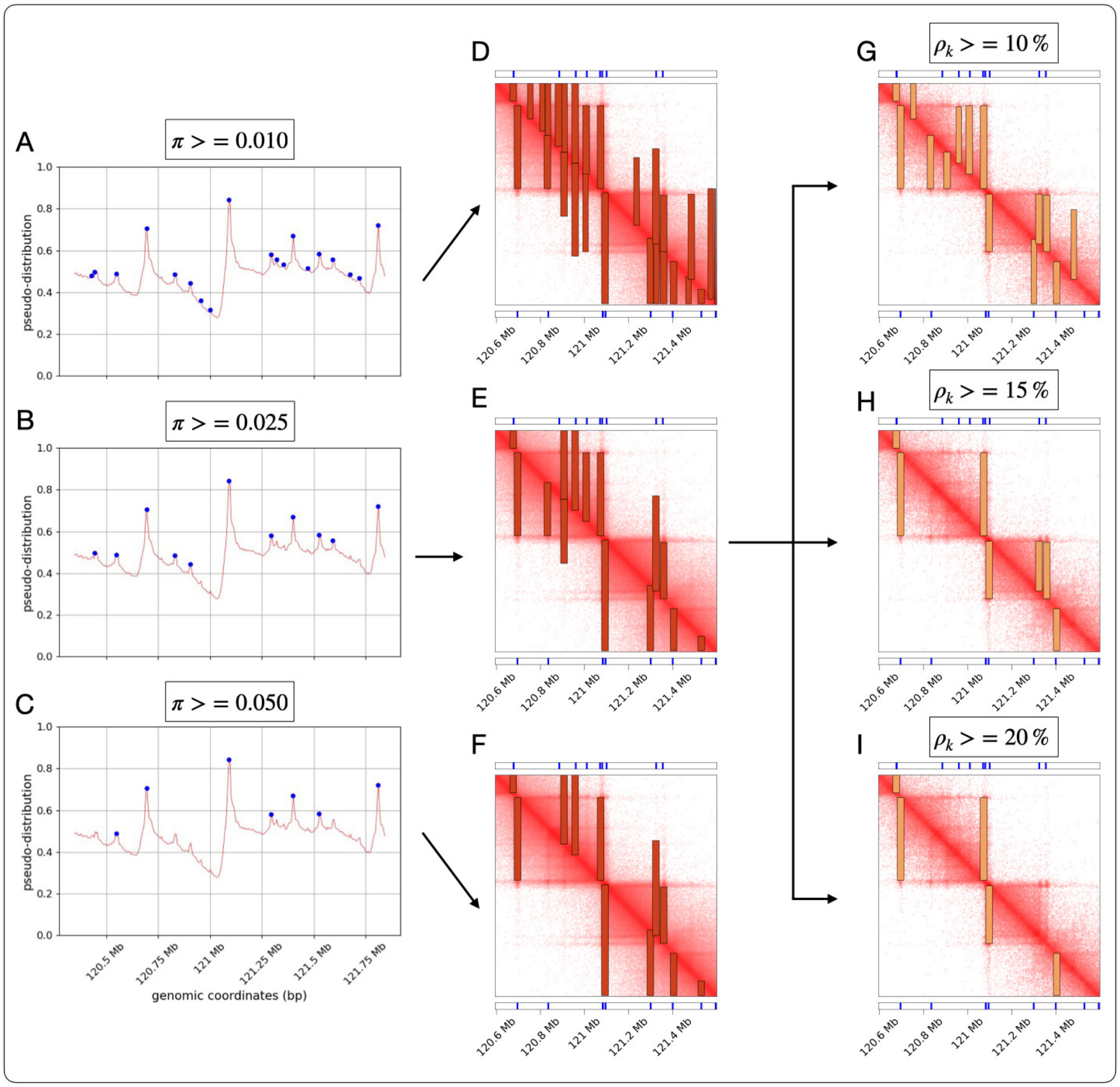


**Supplementary Fig. 8. | StripePy's persistence vs. relative change filterings**. Topological persistence rules out local maximum points (**A**, **B**, and **C**) in terms of the strength of the linear pattern that each bin exhibit. Three thresholds therefore produce as many collections of candidate stripes (**D**, **E**, and **F**) at the end of Step 3. In Step 4, thresholding relative changes further restricts the number of predicted stripes, this time on the basis of a metric that takes into account a well-defined rectangular region (**G**, **H**, and **I**).

Supplementary Table 1. Number of predicted stripes in the StripeBench benchmark. The following encoding is adopted: M1 = StripePy, M2=Chromosight, M3=StripeCaller, M4=Stripenn.

| | | $\sigma = 0kb$ | | | | $\sigma = 5kb$ | | | | $\sigma = 10kb$ | | | | $\sigma = 15kb$ | | | |
|---|---|---|---|---|---|---|---|---|---|---|---|---|---|---|---|---|---|
| $\rho$ | $\delta$ | M1 | M2 | M3 | M4 | M1 | M2 | M3 | M4 | M1 | M2 | M3 | M4 | M1 | M2 | M3 | M4 |
| 5kb | 1 | 35,068 | 30,811 | 19,877 | 274 | 29,868 | 19,143 | 12,103 | 85 | 24,660 | 14,666 | 4,792 | 37 | 19,579 | 10,401 | 731 | 1 |
| | 5 | 31,340 | 33,949 | 29,251 | 1,408 | 26,979 | 22,190 | 19,948 | 139 | 21,241 | 15,730 | 13,577 | 65 | 15,732 | 12,780 | 6,390 | 16 |
| | 10 | 29,647 | 34,619 | 31,807 | 2,410 | 26,478 | 23,411 | 22,229 | 181 | 20,830 | 16,029 | 16,163 | 84 | 15,292 | 12,465 | 9,501 | 21 |
| | 15 | 28,629 | 34,912 | 33,002 | 3,092 | 26,290 | 23,992 | 23,612 | 221 | 20,565 | 16,458 | 17,417 | 110 | 15,085 | 12,414 | 10,997 | 22 |
| 10kb | 1 | 25,112 | 21,197 | 13,662 | 0 | 23,277 | 16,545 | 9,193 | 0 | 20,504 | 13,204 | 5,372 | 0 | 17,553 | 10,887 | 2,357 | 0 |
| | 5 | 22,266 | 22,238 | 19,749 | 0 | 21,622 | 17,428 | 14,683 | 0 | 18,991 | 14,121 | 10,965 | 0 | 15,919 | 11,528 | 7,570 | 0 |
| | 10 | 21,307 | 29,460 | 0 | 0 | 21,046 | 17,760 | 16,441 | 0 | 18,629 | 14,360 | 12,638 | 0 | 15,567 | 11,806 | 9,297 | 0 |
| | 15 | 20,896 | 22,708 | 23,214 | 1 | 20,786 | 17,912 | 17,359 | 0 | 18,437 | 14,503 | 13,477 | 0 | 15,249 | 11,959 | 10,066 | 0 |
| 25kb | 1 | 16,583 | 11,133 | 1,691 | 0 | 16,181 | 10,150 | 1,078 | 0 | 15,411 | 9,070 | 527 | 0 | 14,104 | 8,036 | 195 | 0 |
| | 5 | 15,188 | 11,291 | 4,050 | 0 | 14,964 | 10,267 | 2,967 | 0 | 14,460 | 9,341 | 1,804 | 0 | 13,169 | 8,250 | 927 | 0 |
| | 10 | 14,680 | 11,327 | 4,963 | 0 | 14,621 | 10,288 | 3,804 | 0 | 14,127 | 9,361 | 2,549 | 0 | 12,838 | 8,335 | 1,431 | 0 |
| | 15 | 14,349 | 11,310 | 5,518 | 0 | 14,433 | 10,331 | 4,292 | 0 | 13,961 | 9,373 | 2,910 | 0 | 12,684 | 8,409 | 1,732 | 0 |
| 50kb | 1 | 9,519 | 3,880 | 1 | 0 | 9,370 | 3,793 | 1 | 0 | 9,331 | 3,633 | 0 | 0 | 9,031 | 3,500 | 0 | 0 |
| | 5 | 8,387 | 5,367 | 11 | 0 | 8,298 | 5,156 | 8 | 0 | 8,337 | 4,817 | 10 | 0 | 8,212 | 4,541 | 3 | 0 |
| | 10 | 8,018 | 5,437 | 23 | 0 | 7,984 | 5,195 | 13 | 0 | 7,982 | 4,863 | 11 | 0 | 7,875 | 4,554 | 7 | 0 |
| | 15 | 7,763 | 5,441 | 26 | 0 | 7,758 | 5,169 | 20 | 0 | 7,758 | 4,866 | 19 | 0 | 7,705 | 4,550 | 13 | 0 |

Supplementary Table 2. Classification and recognition measures for each Hi-C matrix in the StripeBench benchmark – resolution set to 5 kb. The following encoding is adopted: M1 = StripePy, M2=Chromosight, M3=StripeCaller, M4=Stripenn.

| | | $\delta = 1$ | | | | $\delta = 5$ | | | | $\delta = 10$ | | | | $\delta = 15$ | | | |
|---|---|---|---|---|---|---|---|---|---|---|---|---|---|---|---|---|---|
| **$\sigma$** | **measure** | M1 | M2 | M3 | M4 | M1 | M2 | M3 | M4 | M1 | M2 | M3 | M4 | M1 | M2 | M3 | M4 |
| 0k | TPR | 36.13 | 34.64 | 28.03 | 0.51 | 35.33 | 35.02 | 33.53 | 1.72 | 34.54 | 35.12 | 33.90 | 2.81 | 33.95 | 35.16 | 33.99 | 3.51 |
| | TNR | 98.03 | 98.34 | 99.08 | 99.99 | 98.32 | 98.09 | 98.45 | 99.93 | 98.44 | 98.04 | 98.24 | 99.87 | 98.51 | 98.02 | 98.15 | 99.83 |
| | PPV | 32.37 | 35.33 | 44.31 | 58.03 | 35.42 | 32.42 | 36.02 | 38.42 | 36.61 | 31.88 | 33.49 | 36.64 | 37.26 | 31.65 | 32.36 | 35.64 |
| | bACC | 67.08 | 66.49 | 63.56 | 50.25 | 66.82 | 66.56 | 65.99 | 50.82 | 66.49 | 66.58 | 66.07 | 51.34 | 66.23 | 66.59 | 66.07 | 51.67 |
| | GM | 59.51 | 58.37 | 52.70 | 7.11 | 58.94 | 58.61 | 57.45 | 13.12 | 58.31 | 58.68 | 57.71 | 16.75 | 57.83 | 58.71 | 57.76 | 18.71 |
| | JI | 20.59 | 21.20 | 20.73 | 0.50 | 21.49 | 20.24 | 21.01 | 1.68 | 21.62 | 20.06 | 20.26 | 2.68 | 21.60 | 19.99 | 19.87 | 3.30 |
| | F1c | 34.15 | 34.98 | 34.34 | 1.00 | 35.38 | 33.67 | 34.73 | 3.30 | 35.55 | 33.42 | 33.69 | 5.22 | 35.53 | 33.31 | 33.15 | 6.39 |
| | FMc | 34.20 | 34.99 | 35.24 | 5.42 | 35.38 | 33.69 | 34.75 | 8.13 | 35.56 | 33.46 | 33.69 | 10.15 | 35.56 | 33.36 | 33.16 | 11.18 |
| | AHR | 83.58 | 34.64 | 28.03 | 1.09 | 81.13 | 35.02 | 33.53 | 5.34 | 79.45 | 35.12 | 33.90 | 9.06 | 78.07 | 35.16 | 33.99 | 11.52 |
| | FGC | 69.66 | 35.33 | 44.31 | 100.00 | 75.76 | 32.42 | 36.02 | 100.00 | 78.39 | 31.88 | 33.49 | 99.96 | 79.73 | 31.65 | 32.36 | 100.00 |
| | F1r | 75.99 | 34.98 | 34.34 | 2.15 | 78.35 | 33.67 | 34.73 | 10.14 | 78.92 | 33.42 | 33.69 | 16.62 | 78.89 | 33.31 | 33.15 | 20.66 |
| | FMr | 76.30 | 34.99 | 35.24 | 10.42 | 78.39 | 33.69 | 34.75 | 23.12 | 78.92 | 33.46 | 33.69 | 30.10 | 78.90 | 33.36 | 33.16 | 33.94 |
| 5k | TPR | 35.91 | 27.59 | 18.13 | 0.14 | 35.82 | 29.53 | 28.78 | 0.25 | 35.53 | 29.78 | 30.65 | 0.30 | 35.57 | 29.91 | 31.62 | 0.43 |
| | TNR | 98.46 | 99.13 | 99.47 | 100.00 | 98.69 | 98.93 | 99.09 | 100.00 | 98.73 | 98.83 | 98.95 | 99.99 | 98.74 | 98.79 | 98.86 | 99.99 |
| | PPV | 37.78 | 45.29 | 47.08 | 52.94 | 41.72 | 41.81 | 45.33 | 56.83 | 42.16 | 39.98 | 43.32 | 52.49 | 42.52 | 39.17 | 42.08 | 61.54 |
| | bACC | 67.18 | 63.36 | 58.80 | 50.07 | 67.26 | 64.23 | 63.93 | 50.12 | 67.13 | 64.31 | 64.80 | 50.15 | 67.16 | 64.35 | 65.24 | 50.21 |
| | GM | 59.46 | 52.30 | 42.47 | 3.78 | 59.46 | 54.05 | 53.40 | 5.01 | 59.23 | 54.26 | 55.07 | 5.50 | 59.27 | 54.36 | 55.91 | 6.58 |
| | JI | 22.56 | 20.69 | 15.06 | 0.14 | 23.87 | 20.93 | 21.36 | 0.25 | 23.89 | 20.58 | 21.88 | 0.30 | 24.02 | 20.42 | 22.03 | 0.43 |
| | F1c | 36.82 | 34.29 | 26.18 | 0.29 | 38.55 | 34.61 | 35.20 | 0.50 | 38.56 | 34.14 | 35.90 | 0.60 | 38.74 | 33.92 | 36.10 | 0.86 |
| | FMc | 36.83 | 35.35 | 29.22 | 2.75 | 38.66 | 35.14 | 36.12 | 3.78 | 38.70 | 34.51 | 36.44 | 3.98 | 38.89 | 34.23 | 36.47 | 5.16 |
| | AHR | 85.60 | 27.59 | 18.13 | 0.36 | 85.54 | 29.53 | 28.78 | 0.57 | 85.30 | 29.78 | 30.65 | 0.76 | 85.20 | 29.91 | 31.62 | 0.94 |
| | FGC | 78.92 | 45.29 | 47.08 | 100.00 | 86.90 | 41.81 | 45.33 | 100.00 | 88.24 | 39.98 | 43.32 | 100.00 | 88.71 | 39.17 | 42.08 | 100.00 |
| | F1r | 82.12 | 34.29 | 26.18 | 0.72 | 86.22 | 34.61 | 35.20 | 1.13 | 86.74 | 34.14 | 35.90 | 1.51 | 86.92 | 33.92 | 36.10 | 1.85 |
| | FMr | 82.19 | 35.35 | 29.22 | 6.00 | 86.22 | 35.14 | 36.12 | 7.55 | 86.76 | 34.51 | 36.44 | 8.72 | 86.94 | 34.23 | 36.47 | 9.67 |
| 10k | TPR | 26.72 | 16.09 | 6.71 | 0.05 | 29.18 | 19.78 | 18.94 | 0.07 | 29.20 | 20.45 | 22.38 | 0.09 | 29.22 | 21.14 | 24.35 | 0.11 |
| | TNR | 98.65 | 99.20 | 99.78 | 100.00 | 99.00 | 99.21 | 99.37 | 100.00 | 99.03 | 99.20 | 99.24 | 100.00 | 99.05 | 99.18 | 99.19 | 99.99 |
| | PPV | 34.05 | 34.47 | 43.99 | 43.24 | 43.17 | 39.51 | 43.83 | 33.85 | 44.05 | 40.09 | 43.50 | 32.14 | 44.65 | 40.37 | 43.93 | 30.00 |
| | bACC | 62.69 | 57.64 | 53.24 | 50.02 | 64.09 | 59.49 | 59.15 | 50.03 | 64.12 | 59.83 | 60.81 | 50.04 | 64.14 | 60.16 | 61.77 | 50.05 |
| | GM | 51.34 | 39.95 | 25.87 | 2.26 | 53.75 | 44.30 | 43.38 | 2.65 | 53.77 | 45.04 | 47.12 | 2.93 | 53.80 | 45.79 | 49.15 | 3.24 |
| | JI | 17.61 | 12.32 | 6.18 | 0.05 | 21.08 | 15.18 | 15.24 | 0.07 | 21.30 | 15.66 | 17.34 | 0.09 | 21.45 | 16.11 | 18.58 | 0.10 |
| | F1c | 29.94 | 21.94 | 11.64 | 0.10 | 34.82 | 26.36 | 26.45 | 0.14 | 35.12 | 27.08 | 29.55 | 0.17 | 35.33 | 27.75 | 31.33 | 0.21 |
| | FMc | 30.17 | 23.55 | 17.18 | 1.48 | 35.49 | 27.95 | 28.81 | 1.54 | 35.86 | 28.63 | 31.20 | 1.66 | 36.12 | 29.22 | 32.71 | 1.77 |
| | AHR | 76.29 | 16.09 | 6.71 | 0.18 | 75.78 | 19.78 | 18.94 | 0.27 | 75.18 | 20.45 | 22.38 | 0.37 | 74.89 | 21.14 | 24.35 | 0.45 |
| | FGC | 83.81 | 34.47 | 43.99 | 100.00 | 95.57 | 39.51 | 43.83 | 100.00 | 96.38 | 40.09 | 43.50 | 100.00 | 97.11 | 40.37 | 43.93 | 100.00 |
| | F1r | 79.87 | 21.94 | 11.64 | 0.36 | 84.53 | 26.36 | 26.45 | 0.53 | 84.47 | 27.08 | 29.55 | 0.74 | 84.57 | 27.75 | 31.33 | 0.89 |
| | FMr | 79.96 | 23.55 | 17.18 | 4.22 | 85.10 | 27.95 | 28.81 | 5.17 | 85.12 | 28.63 | 31.20 | 6.10 | 85.28 | 29.22 | 32.71 | 6.70 |
| 15k | TPR | 16.06 | 7.86 | 0.84 | 0.00 | 18.53 | 10.83 | 7.45 | 0.02 | 19.11 | 11.47 | 11.24 | 0.03 | 19.51 | 11.82 | 13.09 | 0.03 |
| | TNR | 98.79 | 99.34 | 99.96 | 100.00 | 99.18 | 99.22 | 99.66 | 100.00 | 99.23 | 99.26 | 99.50 | 100.00 | 99.26 | 99.28 | 99.43 | 100.00 |
| | PPV | 25.77 | 23.74 | 36.25 | 100.00 | 37.02 | 26.64 | 36.64 | 37.50 | 39.28 | 28.91 | 37.19 | 42.86 | 40.64 | 29.93 | 37.39 | 45.45 |
| | bACC | 57.43 | 53.60 | 50.40 | 50.00 | 58.86 | 55.03 | 53.56 | 50.01 | 59.17 | 55.37 | 55.37 | 50.01 | 59.38 | 55.55 | 56.26 | 50.02 |
| | GM | 39.83 | 27.94 | 9.18 | 0.56 | 42.87 | 32.78 | 27.25 | 1.38 | 43.55 | 33.74 | 33.45 | 1.69 | 44.00 | 34.26 | 36.07 | 1.78 |
| | JI | 10.98 | 6.27 | 0.83 | 0.00 | 14.09 | 8.34 | 6.60 | 0.02 | 14.75 | 8.95 | 9.45 | 0.03 | 15.18 | 9.26 | 10.73 | 0.03 |
| | F1c | 19.79 | 11.81 | 1.65 | 0.01 | 24.70 | 15.40 | 12.38 | 0.04 | 25.71 | 16.42 | 17.27 | 0.06 | 26.36 | 16.95 | 19.39 | 0.06 |
| | FMc | 20.34 | 13.66 | 5.53 | 0.56 | 26.19 | 16.99 | 16.52 | 0.85 | 27.40 | 18.21 | 20.45 | 1.11 | 28.16 | 18.81 | 22.12 | 1.20 |
| | AHR | 60.51 | 7.86 | 0.84 | 0.00 | 59.80 | 10.83 | 7.45 | 0.07 | 58.87 | 11.47 | 11.24 | 0.11 | 58.44 | 11.82 | 13.09 | 0.12 |
| | FGC | 83.29 | 23.74 | 36.25 | 100.00 | 98.94 | 26.64 | 36.64 | 100.00 | 99.61 | 28.91 | 37.19 | 100.00 | 99.72 | 29.93 | 37.39 | 100.00 |
| | F1r | 70.09 | 11.81 | 1.65 | 0.01 | 74.54 | 15.40 | 12.38 | 0.15 | 74.01 | 16.42 | 17.27 | 0.22 | 73.70 | 16.95 | 19.39 | 0.25 |
| | FMr | 70.99 | 13.66 | 5.53 | 0.56 | 76.92 | 16.99 | 16.52 | 2.71 | 76.58 | 18.21 | 20.45 | 3.34 | 76.34 | 18.81 | 22.12 | 3.52 |

Supplementary Table 3. Classification and recognition measures for each Hi-C matrix in the StripeBench benchmark – resolution set to 10 kb. The following encoding is adopted: M1 = StripePy, M2=Chromosight, M3=StripeCaller, M4=Stripenn.

| | | $\delta = 1$ | | | | $\delta = 5$ | | | | $\delta = 10$ | | | | $\delta = 15$ | | | |
|---|---|---|---|---|---|---|---|---|---|---|---|---|---|---|---|---|---|
| $\sigma$ | measure | M1 | M2 | M3 | M4 | M1 | M2 | M3 | M4 | M1 | M2 | M3 | M4 | M1 | M2 | M3 | M4 |
| 0k | TPR | 36.27 | 32.93 | 24.10 | 0.00 | 35.01 | 33.54 | 33.17 | 0.00 | 34.10 | 42.43 | 0.00 | 0.00 | 33.69 | 33.75 | 36.33 | 0.00 |
| | TNR | 97.61 | 98.10 | 98.93 | 100.00 | 98.03 | 97.96 | 98.36 | 100.00 | 98.15 | 97.19 | 100.00 | 100.00 | 98.20 | 97.89 | 97.94 | 100.00 |
| | PPV | 44.17 | 47.51 | 53.95 | 0.00 | 48.08 | 46.13 | 51.36 | 0.00 | 48.94 | 44.04 | 0.00 | 0.00 | 49.30 | 45.45 | 47.86 | 100.00 |
| | bACC | 66.94 | 65.52 | 61.52 | 50.00 | 66.52 | 65.75 | 65.77 | 50.00 | 66.12 | 69.81 | 50.00 | 50.00 | 65.94 | 65.82 | 67.14 | 50.00 |
| | GM | 59.50 | 56.84 | 48.83 | 0.00 | 58.58 | 57.32 | 57.12 | 0.00 | 57.85 | 64.22 | 0.00 | 0.00 | 57.51 | 57.48 | 59.65 | 0.57 |
| | JI | 24.87 | 24.14 | 19.99 | 0.00 | 25.40 | 24.10 | 25.24 | 0.00 | 25.15 | 27.57 | 0.00 | 0.00 | 25.02 | 24.02 | 26.03 | 0.00 |
| | F1c | 39.84 | 38.90 | 33.32 | 0.00 | 40.51 | 38.84 | 40.31 | 0.00 | 40.19 | 43.22 | 0.00 | 0.00 | 40.03 | 38.73 | 41.31 | 0.01 |
| | FMc | 40.03 | 39.55 | 36.06 | 0.00 | 41.02 | 39.34 | 41.27 | 0.00 | 40.85 | 43.23 | 0.00 | 0.00 | 40.75 | 39.16 | 41.70 | 0.57 |
| | AHR | 77.56 | 32.93 | 24.10 | 0.00 | 74.57 | 33.54 | 33.17 | 0.00 | 72.98 | 42.43 | 0.00 | 0.00 | 72.33 | 33.75 | 36.33 | 0.01 |
| | FGC | 83.39 | 47.51 | 53.95 | 0.00 | 89.91 | 46.13 | 51.36 | 0.00 | 91.73 | 44.04 | 0.00 | 0.00 | 92.73 | 45.45 | 47.86 | 100.00 |
| | F1r | 80.37 | 38.90 | 33.32 | 0.00 | 81.53 | 38.84 | 40.31 | 0.00 | 81.28 | 43.22 | 0.00 | 0.00 | 81.27 | 38.73 | 41.31 | 0.01 |
| | FMr | 80.42 | 39.55 | 36.06 | 0.00 | 81.88 | 39.34 | 41.27 | 0.00 | 81.82 | 43.23 | 0.00 | 0.00 | 81.90 | 39.16 | 41.70 | 0.81 |
| 5k | TPR | 36.49 | 28.81 | 16.75 | 0.00 | 36.27 | 29.39 | 25.81 | 0.00 | 35.76 | 29.57 | 28.74 | 0.00 | 35.67 | 29.57 | 30.09 | 0.00 |
| | TNR | 97.94 | 98.68 | 99.31 | 100.00 | 98.21 | 98.56 | 98.84 | 100.00 | 98.28 | 98.52 | 98.70 | 100.00 | 98.32 | 98.49 | 98.61 | 100.00 |
| | PPV | 47.94 | 53.25 | 55.71 | 0.00 | 51.30 | 51.58 | 53.76 | 0.00 | 51.96 | 50.92 | 53.46 | 0.00 | 52.47 | 50.49 | 53.01 | 0.00 |
| | bACC | 67.21 | 63.75 | 58.03 | 50.00 | 67.24 | 63.98 | 62.33 | 50.00 | 67.02 | 64.04 | 63.72 | 50.00 | 66.99 | 64.03 | 64.35 | 50.00 |
| | GM | 59.78 | 53.32 | 40.78 | 0.00 | 59.69 | 53.83 | 50.51 | 0.00 | 59.28 | 53.97 | 53.26 | 0.00 | 59.22 | 53.97 | 54.47 | 0.00 |
| | JI | 26.13 | 23.00 | 14.78 | 0.00 | 26.98 | 23.04 | 21.12 | 0.00 | 26.87 | 23.01 | 22.99 | 0.00 | 26.96 | 22.92 | 23.75 | 0.00 |
| | F1c | 41.44 | 37.39 | 25.75 | 0.00 | 42.50 | 37.45 | 34.88 | 0.00 | 42.36 | 37.41 | 37.39 | 0.00 | 42.47 | 37.30 | 38.39 | 0.00 |
| | FMc | 41.82 | 39.17 | 30.54 | 0.00 | 43.14 | 38.94 | 37.25 | 0.00 | 43.10 | 38.80 | 39.20 | 0.00 | 43.26 | 38.64 | 39.94 | 0.00 |
| | AHR | 79.38 | 28.81 | 16.75 | 0.00 | 78.44 | 29.39 | 25.81 | 0.00 | 77.74 | 29.57 | 28.74 | 0.00 | 77.42 | 29.57 | 30.09 | 0.00 |
| | FGC | 88.47 | 53.25 | 55.71 | 0.00 | 93.65 | 51.58 | 53.76 | 0.00 | 95.29 | 50.92 | 53.46 | 0.00 | 95.97 | 50.49 | 53.01 | 0.00 |
| | F1r | 83.67 | 37.39 | 25.75 | 0.00 | 85.38 | 37.45 | 34.88 | 0.00 | 85.63 | 37.41 | 37.39 | 0.00 | 85.71 | 37.30 | 38.39 | 0.00 |
| | FMr | 83.80 | 39.17 | 30.54 | 0.00 | 85.71 | 38.94 | 37.25 | 0.00 | 86.07 | 38.80 | 39.20 | 0.00 | 86.20 | 38.64 | 39.94 | 0.00 |
| 10k | TPR | 33.26 | 22.91 | 9.85 | 0.00 | 32.86 | 24.81 | 19.32 | 0.00 | 32.74 | 25.25 | 22.25 | 0.00 | 32.57 | 25.52 | 23.48 | 0.00 |
| | TNR | 98.24 | 98.94 | 99.60 | 100.00 | 98.48 | 98.89 | 99.14 | 100.00 | 98.53 | 98.87 | 99.01 | 100.00 | 98.56 | 98.86 | 98.93 | 100.00 |
| | PPV | 49.61 | 53.07 | 56.07 | 0.00 | 52.91 | 53.73 | 53.88 | 0.00 | 53.74 | 53.78 | 53.83 | 0.00 | 54.02 | 53.81 | 53.27 | 0.00 |
| | bACC | 65.75 | 60.93 | 54.72 | 50.00 | 65.67 | 61.85 | 59.23 | 50.00 | 65.64 | 62.06 | 60.63 | 50.00 | 65.56 | 62.19 | 61.20 | 50.00 |
| | GM | 57.16 | 47.61 | 31.32 | 0.00 | 56.89 | 49.53 | 43.76 | 0.00 | 56.80 | 49.97 | 46.93 | 0.00 | 56.66 | 50.23 | 48.19 | 0.00 |
| | JI | 24.86 | 19.05 | 9.14 | 0.00 | 25.43 | 20.44 | 16.58 | 0.00 | 25.54 | 20.75 | 18.68 | 0.00 | 25.50 | 20.93 | 19.47 | 0.00 |
| | F1c | 39.82 | 32.01 | 16.76 | 0.00 | 40.54 | 33.94 | 28.44 | 0.00 | 40.69 | 34.37 | 31.48 | 0.00 | 40.64 | 34.62 | 32.59 | 0.00 |
| | FMc | 40.62 | 34.87 | 23.50 | 0.00 | 41.70 | 36.51 | 32.26 | 0.00 | 41.95 | 36.85 | 34.60 | 0.00 | 41.95 | 37.06 | 35.36 | 0.00 |
| | AHR | 74.63 | 22.91 | 9.85 | 0.00 | 73.19 | 24.81 | 19.32 | 0.00 | 72.45 | 25.25 | 22.25 | 0.00 | 72.05 | 25.52 | 23.48 | 0.00 |
| | FGC | 92.29 | 53.07 | 56.07 | 0.00 | 97.30 | 53.73 | 53.88 | 0.00 | 98.04 | 53.78 | 53.83 | 0.00 | 98.47 | 53.81 | 53.27 | 0.00 |
| | F1r | 82.52 | 32.01 | 16.76 | 0.00 | 83.54 | 33.94 | 28.44 | 0.00 | 83.33 | 34.37 | 31.48 | 0.00 | 83.21 | 34.62 | 32.59 | 0.00 |
| | FMr | 82.99 | 34.87 | 23.50 | 0.00 | 84.39 | 36.51 | 32.26 | 0.00 | 84.28 | 36.85 | 34.60 | 0.00 | 84.23 | 37.06 | 35.36 | 0.00 |
| 15k | TPR | 27.19 | 16.46 | 4.04 | 0.00 | 27.24 | 18.62 | 13.03 | 0.00 | 26.79 | 19.65 | 16.04 | 0.00 | 26.37 | 20.01 | 17.37 | 0.00 |
| | TNR | 98.43 | 99.00 | 99.81 | 100.00 | 98.71 | 99.01 | 99.39 | 100.00 | 98.74 | 99.01 | 99.25 | 100.00 | 98.78 | 99.01 | 99.19 | 100.00 |
| | PPV | 47.37 | 46.24 | 52.48 | 0.00 | 52.32 | 49.39 | 52.64 | 0.00 | 52.63 | 50.90 | 52.77 | 0.00 | 52.88 | 51.16 | 52.78 | 0.00 |
| | bACC | 62.81 | 57.73 | 51.93 | 50.00 | 62.97 | 58.81 | 56.21 | 50.00 | 62.77 | 59.33 | 57.65 | 50.00 | 62.57 | 59.51 | 58.28 | 50.00 |
| | GM | 51.73 | 40.37 | 20.09 | 0.00 | 51.85 | 42.94 | 35.99 | 0.00 | 51.43 | 44.11 | 39.90 | 0.00 | 51.03 | 44.50 | 41.51 | 0.00 |
| | JI | 20.88 | 13.82 | 3.90 | 0.00 | 21.82 | 15.64 | 11.66 | 0.00 | 21.59 | 16.52 | 14.03 | 0.00 | 21.35 | 16.80 | 15.04 | 0.00 |
| | F1c | 34.55 | 24.28 | 7.51 | 0.00 | 35.82 | 27.04 | 20.89 | 0.00 | 35.51 | 28.35 | 24.61 | 0.00 | 35.19 | 28.76 | 26.14 | 0.00 |
| | FMc | 35.89 | 27.59 | 14.57 | 0.00 | 37.75 | 30.33 | 26.19 | 0.00 | 37.55 | 31.62 | 29.10 | 0.00 | 37.34 | 31.99 | 30.28 | 0.00 |
| | AHR | 67.04 | 16.46 | 4.04 | 0.00 | 63.98 | 18.62 | 13.03 | 0.00 | 63.17 | 19.65 | 16.04 | 0.00 | 62.19 | 20.01 | 17.37 | 0.00 |
| | FGC | 94.91 | 46.24 | 52.48 | 0.00 | 99.05 | 49.39 | 52.64 | 0.00 | 99.43 | 50.90 | 52.77 | 0.00 | 99.61 | 51.16 | 52.78 | 0.00 |
| | F1r | 78.57 | 24.28 | 7.51 | 0.00 | 77.74 | 27.04 | 20.89 | 0.00 | 77.25 | 28.35 | 24.61 | 0.00 | 76.57 | 28.76 | 26.14 | 0.00 |
| | FMr | 79.76 | 27.59 | 14.57 | 0.00 | 79.60 | 30.33 | 26.19 | 0.00 | 79.25 | 31.62 | 29.10 | 0.00 | 78.71 | 31.99 | 30.28 | 0.00 |

Supplementary Table 4. Classification and recognition measures for each Hi-C matrix in the StripeBench benchmark – resolution set to 25 kb. The following encoding is adopted: M1 = StripePy, M2=Chromosight, M3=StripeCaller, M4=Stripenn.

| | | $\delta = 1$ | | | | $\delta = 5$ | | | | $\delta = 10$ | | | | $\delta = 15$ | | | |
|---|---|---|---|---|---|---|---|---|---|---|---|---|---|---|---|---|---|
| $\sigma$ | **measure** | M1 | M2 | M3 | M4 | M1 | M2 | M3 | M4 | M1 | M2 | M3 | M4 | M1 | M2 | M3 | M4 |
| 0k | TPR | 29.89 | 21.04 | 3.30 | 0.00 | 28.81 | 21.49 | 7.98 | 0.00 | 27.99 | 21.54 | 9.66 | 0.00 | 27.52 | 21.46 | 10.80 | 0.00 |
| | TNR | 96.33 | 97.66 | 99.66 | 100.00 | 96.82 | 97.65 | 99.19 | 100.00 | 96.95 | 97.64 | 98.99 | 100.00 | 97.04 | 97.64 | 98.89 | 100.00 |
| | PPV | 51.61 | 54.10 | 55.82 | 0.00 | 54.31 | 54.50 | 56.44 | 0.00 | 54.60 | 54.44 | 55.73 | 0.00 | 54.90 | 54.33 | 56.03 | 0.00 |
| | bACC | 63.11 | 59.35 | 51.48 | 50.00 | 62.82 | 59.57 | 53.59 | 50.00 | 62.47 | 59.59 | 54.33 | 50.00 | 62.28 | 59.55 | 54.84 | 50.00 |
| | GM | 53.66 | 45.33 | 18.13 | 0.00 | 52.81 | 45.81 | 28.14 | 0.00 | 52.10 | 45.86 | 30.93 | 0.00 | 51.67 | 45.78 | 32.68 | 0.00 |
| | JI | 23.35 | 17.85 | 3.21 | 0.00 | 23.19 | 18.22 | 7.52 | 0.00 | 22.71 | 18.25 | 8.97 | 0.00 | 22.44 | 18.18 | 9.96 | 0.00 |
| | F1c | 37.86 | 30.29 | 6.23 | 0.00 | 37.65 | 30.83 | 13.99 | 0.00 | 37.01 | 30.86 | 16.47 | 0.00 | 36.66 | 30.77 | 18.11 | 0.00 |
| | FMc | 39.28 | 33.74 | 13.57 | 0.00 | 39.55 | 34.23 | 21.23 | 0.00 | 39.10 | 34.24 | 23.20 | 0.00 | 38.87 | 34.15 | 24.60 | 0.00 |
| | AHR | 66.28 | 21.04 | 3.30 | 0.00 | 63.85 | 21.49 | 7.98 | 0.00 | 62.32 | 21.54 | 9.66 | 0.00 | 61.36 | 21.46 | 10.80 | 0.00 |
| | FGC | 94.28 | 54.10 | 55.82 | 0.00 | 98.29 | 54.50 | 56.44 | 0.00 | 98.86 | 54.44 | 55.73 | 0.00 | 99.16 | 54.33 | 56.03 | 0.00 |
| | F1r | 77.84 | 30.29 | 6.23 | 0.00 | 77.41 | 30.83 | 13.99 | 0.00 | 76.44 | 30.86 | 16.47 | 0.00 | 75.81 | 30.77 | 18.11 | 0.00 |
| | FMr | 79.05 | 33.74 | 13.57 | 0.00 | 79.22 | 34.23 | 21.23 | 0.00 | 78.49 | 34.24 | 23.20 | 0.00 | 78.00 | 34.15 | 24.60 | 0.00 |
| 5k | TPR | 29.99 | 19.49 | 2.22 | 0.00 | 28.80 | 19.89 | 5.96 | 0.00 | 28.46 | 19.91 | 7.61 | 0.00 | 28.15 | 20.01 | 8.52 | 0.00 |
| | TNR | 96.52 | 97.91 | 99.80 | 100.00 | 96.92 | 97.91 | 99.42 | 100.00 | 97.04 | 97.90 | 99.26 | 100.00 | 97.08 | 97.89 | 99.15 | 100.00 |
| | PPV | 53.07 | 54.97 | 59.00 | 0.00 | 55.10 | 55.47 | 57.47 | 0.00 | 55.72 | 55.40 | 57.26 | 0.00 | 55.84 | 55.45 | 56.85 | 0.00 |
| | bACC | 63.26 | 58.70 | 51.01 | 50.00 | 62.86 | 58.90 | 52.69 | 50.00 | 62.75 | 58.90 | 53.43 | 50.00 | 62.61 | 58.95 | 53.84 | 50.00 |
| | GM | 53.80 | 43.68 | 14.89 | 0.00 | 52.83 | 44.13 | 24.33 | 0.00 | 52.55 | 44.15 | 27.48 | 0.00 | 52.27 | 44.26 | 29.07 | 0.00 |
| | JI | 23.70 | 16.80 | 2.19 | 0.00 | 23.32 | 17.15 | 5.70 | 0.00 | 23.21 | 17.16 | 7.20 | 0.00 | 23.02 | 17.24 | 8.00 | 0.00 |
| | F1c | 38.32 | 28.77 | 4.28 | 0.00 | 37.83 | 29.28 | 10.79 | 0.00 | 37.67 | 29.29 | 13.43 | 0.00 | 37.43 | 29.41 | 14.82 | 0.00 |
| | FMc | 39.90 | 32.73 | 11.45 | 0.00 | 39.83 | 33.22 | 18.50 | 0.00 | 39.82 | 33.21 | 20.87 | 0.00 | 39.64 | 33.31 | 22.01 | 0.00 |
| | AHR | 66.29 | 19.49 | 2.22 | 0.00 | 64.07 | 19.89 | 5.96 | 0.00 | 63.08 | 19.91 | 7.61 | 0.00 | 62.53 | 20.01 | 8.52 | 0.00 |
| | FGC | 95.33 | 54.97 | 59.00 | 0.00 | 98.80 | 55.47 | 57.47 | 0.00 | 99.26 | 55.40 | 57.26 | 0.00 | 99.43 | 55.45 | 56.85 | 0.00 |
| | F1r | 78.20 | 28.77 | 4.28 | 0.00 | 77.73 | 29.28 | 10.79 | 0.00 | 77.14 | 29.29 | 13.43 | 0.00 | 76.78 | 29.41 | 14.82 | 0.00 |
| | FMr | 79.50 | 32.73 | 11.45 | 0.00 | 79.56 | 33.22 | 18.50 | 0.00 | 79.13 | 33.21 | 20.87 | 0.00 | 78.85 | 33.31 | 22.01 | 0.00 |
| 10k | TPR | 29.27 | 17.58 | 1.11 | 0.00 | 28.24 | 18.29 | 3.77 | 0.00 | 27.89 | 18.38 | 5.34 | 0.00 | 27.64 | 18.34 | 6.02 | 0.00 |
| | TNR | 96.78 | 98.15 | 99.90 | 100.00 | 97.08 | 98.12 | 99.67 | 100.00 | 97.19 | 98.12 | 99.53 | 100.00 | 97.23 | 98.11 | 99.46 | 100.00 |
| | PPV | 54.37 | 55.49 | 60.34 | 0.00 | 55.91 | 56.08 | 59.87 | 0.00 | 56.53 | 56.22 | 59.95 | 0.00 | 56.69 | 56.01 | 59.21 | 0.00 |
| | bACC | 63.02 | 57.87 | 50.51 | 50.00 | 62.66 | 58.21 | 51.72 | 50.00 | 62.54 | 58.25 | 52.43 | 50.00 | 62.44 | 58.22 | 52.74 | 50.00 |
| | GM | 53.22 | 41.54 | 10.53 | 0.00 | 52.36 | 42.37 | 19.39 | 0.00 | 52.07 | 42.47 | 23.05 | 0.00 | 51.85 | 42.42 | 24.46 | 0.00 |
| | JI | 23.49 | 15.41 | 1.10 | 0.00 | 23.09 | 16.00 | 3.68 | 0.00 | 22.97 | 16.08 | 5.15 | 0.00 | 22.82 | 16.03 | 5.78 | 0.00 |
| | F1c | 38.05 | 26.70 | 2.18 | 0.00 | 37.52 | 27.59 | 7.10 | 0.00 | 37.35 | 27.71 | 9.80 | 0.00 | 37.17 | 27.63 | 10.93 | 0.00 |
| | FMc | 39.89 | 31.23 | 8.19 | 0.00 | 39.73 | 32.03 | 15.03 | 0.00 | 39.71 | 32.15 | 17.89 | 0.00 | 39.59 | 32.05 | 18.88 | 0.00 |
| | AHR | 64.49 | 17.58 | 1.11 | 0.00 | 62.75 | 18.29 | 3.77 | 0.00 | 61.63 | 18.38 | 5.34 | 0.00 | 61.21 | 18.34 | 6.02 | 0.00 |
| | FGC | 96.20 | 55.49 | 60.34 | 0.00 | 99.09 | 56.08 | 59.87 | 0.00 | 99.47 | 56.22 | 59.95 | 0.00 | 99.65 | 56.01 | 59.21 | 0.00 |
| | F1r | 77.22 | 26.70 | 2.18 | 0.00 | 76.84 | 27.59 | 7.10 | 0.00 | 76.10 | 27.71 | 9.80 | 0.00 | 75.84 | 27.63 | 10.93 | 0.00 |
| | FMr | 78.77 | 31.23 | 8.19 | 0.00 | 78.85 | 32.03 | 15.03 | 0.00 | 78.29 | 32.15 | 17.89 | 0.00 | 78.10 | 32.05 | 18.88 | 0.00 |
| 15k | TPR | 27.02 | 15.86 | 0.41 | 0.00 | 26.00 | 16.49 | 2.00 | 0.00 | 25.54 | 16.70 | 3.10 | 0.00 | 25.30 | 16.80 | 3.68 | 0.00 |
| | TNR | 97.08 | 98.40 | 99.96 | 100.00 | 97.38 | 98.38 | 99.84 | 100.00 | 97.47 | 98.37 | 99.75 | 100.00 | 97.51 | 98.35 | 99.69 | 100.00 |
| | PPV | 54.85 | 56.51 | 59.49 | 0.00 | 56.53 | 57.21 | 61.70 | 0.00 | 56.96 | 57.35 | 62.05 | 0.00 | 57.10 | 57.21 | 60.91 | 0.00 |
| | bACC | 62.05 | 57.13 | 50.18 | 50.00 | 61.69 | 57.43 | 50.92 | 50.00 | 61.50 | 57.53 | 51.43 | 50.00 | 61.40 | 57.58 | 51.69 | 50.00 |
| | GM | 51.22 | 39.51 | 6.36 | 0.00 | 50.32 | 40.27 | 14.12 | 0.00 | 49.89 | 40.53 | 17.59 | 0.00 | 49.67 | 40.65 | 19.17 | 0.00 |
| | JI | 22.10 | 14.13 | 0.40 | 0.00 | 21.67 | 14.68 | 1.97 | 0.00 | 21.41 | 14.85 | 3.04 | 0.00 | 21.26 | 14.93 | 3.60 | 0.00 |
| | F1c | 36.20 | 24.77 | 0.80 | 0.00 | 35.62 | 25.60 | 3.87 | 0.00 | 35.26 | 25.86 | 5.91 | 0.00 | 35.06 | 25.98 | 6.95 | 0.00 |
| | FMc | 38.50 | 29.94 | 4.91 | 0.00 | 38.34 | 30.71 | 11.10 | 0.00 | 38.14 | 30.94 | 13.87 | 0.00 | 38.01 | 31.01 | 14.98 | 0.00 |
| | AHR | 59.94 | 15.86 | 0.41 | 0.00 | 58.01 | 16.49 | 2.00 | 0.00 | 56.78 | 16.70 | 3.10 | 0.00 | 56.22 | 16.80 | 3.68 | 0.00 |
| | FGC | 96.87 | 56.51 | 59.49 | 0.00 | 99.38 | 57.21 | 61.70 | 0.00 | 99.64 | 57.35 | 62.05 | 0.00 | 99.79 | 57.21 | 60.91 | 0.00 |
| | F1r | 74.06 | 24.77 | 0.80 | 0.00 | 73.26 | 25.60 | 3.87 | 0.00 | 72.34 | 25.86 | 5.91 | 0.00 | 71.92 | 25.98 | 6.95 | 0.00 |
| | FMr | 76.20 | 29.94 | 4.91 | 0.00 | 75.93 | 30.71 | 11.10 | 0.00 | 75.22 | 30.94 | 13.87 | 0.00 | 74.90 | 31.01 | 14.98 | 0.00 |

Supplementary Table 5. Classification and recognition measures for each Hi-C matrix in the StripeBench benchmark – resolution set to 50 kb. The following encoding is adopted: M1 = StripePy, M2=Chromosight, M3=StripeCaller, M4=Stripenn.

| | | $\delta = 1$ | | | | $\delta = 5$ | | | | $\delta = 10$ | | | | $\delta = 15$ | | | |
|---|---|---|---|---|---|---|---|---|---|---|---|---|---|---|---|---|---|
| $\sigma$ | **measure** | M1 | M2 | M3 | M4 | M1 | M2 | M3 | M4 | M1 | M2 | M3 | M4 | M1 | M2 | M3 | M4 |
| 0k | TPR | 21.48 | 8.59 | 0.00 | 0.00 | 19.97 | 12.26 | 0.01 | 0.00 | 19.16 | 12.47 | 0.05 | 0.00 | 18.76 | 12.47 | 0.05 | 0.00 |
| | TNR | 96.00 | 98.32 | 100.00 | 100.00 | 96.76 | 97.79 | 99.99 | 100.00 | 96.92 | 97.77 | 99.99 | 100.00 | 97.07 | 97.77 | 99.99 | 100.00 |
| | PPV | 59.09 | 57.96 | 100.00 | 0.00 | 62.35 | 59.83 | 27.27 | 0.00 | 62.58 | 60.09 | 56.52 | 0.00 | 63.30 | 60.04 | 53.85 | 0.00 |
| | bACC | 58.74 | 53.46 | 50.00 | 50.00 | 58.36 | 55.02 | 50.00 | 50.00 | 58.04 | 55.12 | 50.02 | 50.00 | 57.92 | 55.12 | 50.02 | 50.00 |
| | GM | 45.41 | 29.06 | 0.62 | 0.00 | 43.95 | 34.62 | 1.07 | 0.00 | 43.09 | 34.92 | 2.23 | 0.00 | 42.68 | 34.92 | 2.31 | 0.00 |
| | JI | 18.70 | 8.08 | 0.00 | 0.00 | 17.82 | 11.33 | 0.01 | 0.00 | 17.19 | 11.52 | 0.05 | 0.00 | 16.92 | 11.52 | 0.05 | 0.00 |
| | F1c | 31.50 | 14.96 | 0.01 | 0.00 | 30.25 | 20.35 | 0.02 | 0.00 | 29.34 | 20.66 | 0.10 | 0.00 | 28.95 | 20.66 | 0.11 | 0.00 |
| | FMc | 35.63 | 22.31 | 0.62 | 0.00 | 35.28 | 27.08 | 0.56 | 0.00 | 34.63 | 27.38 | 1.67 | 0.00 | 34.46 | 27.37 | 1.70 | 0.00 |
| | AHR | 49.38 | 8.59 | 0.00 | 0.00 | 46.00 | 12.26 | 0.01 | 0.00 | 44.32 | 12.47 | 0.05 | 0.00 | 43.32 | 12.47 | 0.05 | 0.00 |
| | FGC | 95.85 | 57.96 | 100.00 | 0.00 | 99.24 | 59.83 | 27.27 | 0.00 | 99.35 | 60.09 | 56.52 | 0.00 | 99.65 | 60.04 | 53.85 | 0.00 |
| | F1r | 65.18 | 14.96 | 0.01 | 0.00 | 62.87 | 20.35 | 0.02 | 0.00 | 61.29 | 20.66 | 0.10 | 0.00 | 60.38 | 20.66 | 0.11 | 0.00 |
| | FMr | 68.80 | 22.31 | 0.62 | 0.00 | 67.57 | 27.08 | 0.56 | 0.00 | 66.35 | 27.38 | 1.67 | 0.00 | 65.70 | 27.37 | 1.70 | 0.00 |
| 5k | TPR | 21.60 | 8.39 | 0.00 | 0.00 | 19.99 | 11.82 | 0.02 | 0.00 | 19.48 | 11.95 | 0.03 | 0.00 | 19.00 | 11.99 | 0.05 | 0.00 |
| | TNR | 96.19 | 98.36 | 100.00 | 100.00 | 96.85 | 97.88 | 100.00 | 100.00 | 97.04 | 97.88 | 99.99 | 100.00 | 97.14 | 97.92 | 99.99 | 100.00 |
| | PPV | 60.37 | 57.92 | 0.00 | 0.00 | 63.10 | 60.03 | 62.50 | 0.00 | 63.90 | 60.25 | 53.85 | 0.00 | 64.13 | 60.75 | 70.00 | 0.00 |
| | bACC | 58.89 | 53.37 | 50.00 | 50.00 | 58.42 | 54.85 | 50.01 | 50.00 | 58.26 | 54.92 | 50.01 | 50.00 | 58.07 | 54.95 | 50.02 | 50.00 |
| | GM | 45.58 | 28.72 | 0.00 | 0.00 | 44.00 | 34.01 | 1.38 | 0.00 | 43.48 | 34.20 | 1.63 | 0.00 | 42.96 | 34.26 | 2.31 | 0.00 |
| | JI | 18.92 | 7.91 | 0.00 | 0.00 | 17.90 | 10.96 | 0.02 | 0.00 | 17.55 | 11.08 | 0.03 | 0.00 | 17.17 | 11.13 | 0.05 | 0.00 |
| | F1c | 31.82 | 14.65 | 0.00 | 0.00 | 30.36 | 19.75 | 0.04 | 0.00 | 29.86 | 19.95 | 0.05 | 0.00 | 29.31 | 20.03 | 0.11 | 0.00 |
| | FMc | 36.11 | 22.04 | 0.00 | 0.00 | 35.52 | 26.63 | 1.09 | 0.00 | 35.28 | 26.83 | 1.20 | 0.00 | 34.90 | 26.99 | 1.93 | 0.00 |
| | AHR | 49.31 | 8.39 | 0.00 | 0.00 | 45.98 | 11.82 | 0.02 | 0.00 | 44.66 | 11.95 | 0.03 | 0.00 | 43.68 | 11.99 | 0.05 | 0.00 |
| | FGC | 96.45 | 57.92 | 0.00 | 0.00 | 99.29 | 60.03 | 62.50 | 0.00 | 99.62 | 60.25 | 53.85 | 0.00 | 99.73 | 60.75 | 70.00 | 0.00 |
| | F1r | 65.26 | 14.65 | 0.00 | 0.00 | 62.85 | 19.75 | 0.04 | 0.00 | 61.68 | 19.95 | 0.05 | 0.00 | 60.75 | 20.03 | 0.11 | 0.00 |
| | FMr | 68.97 | 22.04 | 0.00 | 0.00 | 67.57 | 26.63 | 1.09 | 0.00 | 66.71 | 26.83 | 1.20 | 0.00 | 66.00 | 26.99 | 1.93 | 0.00 |
| 10k | TPR | 21.85 | 8.08 | 0.00 | 0.00 | 20.41 | 11.33 | 0.02 | 0.00 | 19.72 | 11.45 | 0.02 | 0.00 | 19.34 | 11.52 | 0.03 | 0.00 |
| | TNR | 96.29 | 98.44 | 100.00 | 100.00 | 96.93 | 98.10 | 99.99 | 100.00 | 97.11 | 98.08 | 99.99 | 100.00 | 97.23 | 98.10 | 99.99 | 100.00 |
| | PPV | 61.33 | 58.24 | 0.00 | 0.00 | 64.12 | 61.62 | 50.00 | 0.00 | 64.71 | 61.65 | 45.45 | 0.00 | 65.27 | 61.98 | 36.84 | 0.00 |
| | bACC | 59.07 | 53.26 | 50.00 | 50.00 | 58.67 | 54.72 | 50.01 | 50.00 | 58.41 | 54.77 | 50.01 | 50.00 | 58.28 | 54.81 | 50.01 | 50.00 |
| | GM | 45.87 | 28.20 | 0.00 | 0.00 | 44.48 | 33.34 | 1.38 | 0.00 | 43.76 | 33.51 | 1.38 | 0.00 | 43.36 | 33.61 | 1.63 | 0.00 |
| | JI | 19.21 | 7.64 | 0.00 | 0.00 | 18.32 | 10.59 | 0.02 | 0.00 | 17.81 | 10.69 | 0.02 | 0.00 | 17.53 | 10.76 | 0.03 | 0.00 |
| | F1c | 32.22 | 14.19 | 0.00 | 0.00 | 30.97 | 19.14 | 0.04 | 0.00 | 30.23 | 19.31 | 0.04 | 0.00 | 29.83 | 19.42 | 0.05 | 0.00 |
| | FMc | 36.61 | 21.69 | 0.00 | 0.00 | 36.18 | 26.42 | 0.98 | 0.00 | 35.72 | 26.57 | 0.93 | 0.00 | 35.53 | 26.72 | 0.99 | 0.00 |
| | AHR | 49.70 | 8.08 | 0.00 | 0.00 | 46.47 | 11.33 | 0.02 | 0.00 | 45.03 | 11.45 | 0.02 | 0.00 | 43.93 | 11.52 | 0.03 | 0.00 |
| | FGC | 96.67 | 58.24 | 0.00 | 0.00 | 99.47 | 61.62 | 50.00 | 0.00 | 99.69 | 61.65 | 45.45 | 0.00 | 99.81 | 61.98 | 36.84 | 0.00 |
| | F1r | 65.65 | 14.19 | 0.00 | 0.00 | 63.35 | 19.14 | 0.04 | 0.00 | 62.04 | 19.31 | 0.04 | 0.00 | 61.01 | 19.42 | 0.05 | 0.00 |
| | FMr | 69.32 | 21.69 | 0.00 | 0.00 | 67.99 | 26.42 | 0.98 | 0.00 | 67.00 | 26.56 | 0.93 | 0.00 | 66.22 | 26.72 | 0.99 | 0.00 |
| 15k | TPR | 21.39 | 7.93 | 0.00 | 0.00 | 20.23 | 10.79 | 0.00 | 0.00 | 19.57 | 10.84 | 0.02 | 0.00 | 19.26 | 10.91 | 0.03 | 0.00 |
| | TNR | 96.48 | 98.54 | 100.00 | 100.00 | 97.01 | 98.24 | 100.00 | 100.00 | 97.18 | 98.24 | 100.00 | 100.00 | 97.27 | 98.26 | 99.99 | 100.00 |
| | PPV | 62.02 | 59.37 | 0.00 | 0.00 | 64.50 | 62.23 | 33.33 | 0.00 | 65.09 | 62.32 | 57.14 | 0.00 | 65.48 | 62.79 | 53.85 | 0.00 |
| | bACC | 58.93 | 53.24 | 50.00 | 50.00 | 58.62 | 54.51 | 50.00 | 50.00 | 58.37 | 54.54 | 50.01 | 50.00 | 58.27 | 54.58 | 50.01 | 50.00 |
| | GM | 45.42 | 27.96 | 0.00 | 0.00 | 44.29 | 32.56 | 0.62 | 0.00 | 43.61 | 32.63 | 1.24 | 0.00 | 43.29 | 32.74 | 1.63 | 0.00 |
| | JI | 18.91 | 7.53 | 0.00 | 0.00 | 18.20 | 10.13 | 0.00 | 0.00 | 17.71 | 10.17 | 0.02 | 0.00 | 17.49 | 10.25 | 0.03 | 0.00 |
| | F1c | 31.80 | 14.00 | 0.00 | 0.00 | 30.79 | 18.39 | 0.01 | 0.00 | 30.10 | 18.46 | 0.03 | 0.00 | 29.77 | 18.59 | 0.05 | 0.00 |
| | FMc | 36.42 | 21.70 | 0.00 | 0.00 | 36.12 | 25.91 | 0.36 | 0.00 | 35.69 | 25.99 | 0.93 | 0.00 | 35.51 | 26.17 | 1.20 | 0.00 |
| | AHR | 48.58 | 7.93 | 0.00 | 0.00 | 46.02 | 10.79 | 0.00 | 0.00 | 44.53 | 10.84 | 0.02 | 0.00 | 43.83 | 10.91 | 0.03 | 0.00 |
| | FGC | 96.78 | 59.37 | 0.00 | 0.00 | 99.55 | 62.23 | 33.33 | 0.00 | 99.77 | 62.32 | 57.14 | 0.00 | 99.86 | 62.79 | 53.85 | 0.00 |
| | F1r | 64.69 | 14.00 | 0.00 | 0.00 | 62.95 | 18.39 | 0.01 | 0.00 | 61.58 | 18.46 | 0.03 | 0.00 | 60.92 | 18.59 | 0.05 | 0.00 |
| | FMr | 68.57 | 21.70 | 0.00 | 0.00 | 67.69 | 25.91 | 0.36 | 0.00 | 66.65 | 25.99 | 0.93 | 0.00 | 66.16 | 26.17 | 1.20 | 0.00 |

Supplementary Table 6. Clamped $p$-values for the Anderson-Darling test run over sub-sets of the StripeBench benchmark (obtained by fixing one between contact density $\delta$, noise level $\sigma$, and resolution $\rho$) and the whole benchmark. The minimum value used for clamping is set to $1e-04$. Values below the significant threshold of 0.05 are written in bold.

| | | methods | TPR | TNR | PPV | bACC | GM | F1C | FMc | JI | AHR | FGC | F1r | FMr |
|---|---|---|---|---|---|---|---|---|---|---|---|---|---|---|
| contact density $\delta$ | 1 | M1, M2 | **2.2e-03** | **1.3e-03** | 6.0e-01 | **6.5e-03** | **2.4e-03** | **3.4e-03** | **3.4e-03** | **5.0e-04** | **1.0e-04** | **1.0e-04** | **1.0e-04** | **1.0e-04** |
| | | M1, M3 | **1.0e-04** | **1.0e-04** | 6.3e-01 | **1.0e-04** | **1.0e-04** | **1.0e-04** | **1.0e-04** | **1.0e-04** | **1.0e-04** | **1.0e-04** | **1.0e-04** | **1.0e-04** |
| | | M2, M3 | **1.1e-03** | **1.0e-04** | 4.2e-01 | **1.4e-03** | **1.1e-03** | **1.0e-03** | **1.0e-03** | **1.6e-03** | **1.1e-03** | 4.2e-01 | **1.0e-03** | **1.6e-03** |
| | 5 | M1, M2 | **7.0e-03** | 5.0e-02 | 6.4e-01 | **3.0e-02** | **1.5e-02** | **1.2e-03** | **1.2e-03** | **3.0e-04** | **1.0e-04** | **1.0e-04** | **1.0e-04** | **1.0e-04** |
| | | M1, M3 | **5.0e-04** | **1.0e-04** | 5.1e-01 | **1.5e-03** | **7.0e-04** | **3.0e-04** | **3.0e-04** | **2.0e-04** | **1.0e-04** | **1.0e-04** | **1.0e-04** | **1.0e-04** |
| | | M2, M3 | **1.8e-02** | **1.0e-04** | 9.6e-01 | **2.3e-02** | **2.0e-02** | **2.1e-02** | **2.1e-02** | **2.1e-02** | **1.8e-02** | 9.6e-01 | **2.1e-02** | **2.1e-02** |
| | 10 | M1, M2 | **4.0e-02** | 8.2e-02 | 5.8e-01 | 5.2e-02 | **4.7e-02** | **2.9e-03** | **2.9e-03** | **4.0e-04** | **1.0e-04** | **1.0e-04** | **1.0e-04** | **1.0e-04** |
| | | M1, M3 | **9.0e-04** | **1.0e-04** | 5.5e-01 | **9.0e-04** | **9.0e-04** | **3.0e-04** | **3.0e-04** | **1.0e-04** | **1.0e-04** | **1.0e-04** | **1.0e-04** | **1.0e-04** |
| | | M2, M3 | **8.5e-03** | **2.0e-04** | 9.2e-01 | **2.0e-02** | **8.5e-03** | **1.0e-02** | **1.0e-02** | **1.4e-02** | **8.5e-03** | 9.2e-01 | **1.0e-02** | **1.4e-02** |
| | 15 | M1, M2 | **4.0e-02** | 6.8e-02 | 5.2e-01 | **4.6e-02** | **3.9e-02** | **2.2e-03** | **2.2e-03** | **2.0e-04** | **1.0e-04** | **1.0e-04** | **1.0e-04** | **1.0e-04** |
| | | M1, M3 | **5.1e-03** | **1.0e-04** | 6.3e-01 | **9.2e-03** | **4.9e-03** | **2.2e-03** | **2.2e-03** | **4.0e-04** | **1.0e-04** | **1.0e-04** | **1.0e-04** | **1.0e-04** |
| | | M2, M3 | 5.3e-02 | **4.0e-04** | 9.2e-01 | 8.6e-02 | 5.7e-02 | **4.8e-02** | **4.8e-02** | **5.0e-02** | 5.3e-02 | 9.2e-01 | **4.8e-02** | **5.0e-02** |
| noise level $\sigma$ | 0 kb | M1, M2 | 4.5e-01 | **3.2e-02** | 6.3e-01 | 4.9e-01 | 4.7e-01 | 7.6e-02 | 7.6e-02 | **1.3e-03** | **1.0e-04** | **1.0e-04** | **1.0e-04** | **1.0e-04** |
| | | M1, M3 | **7.8e-03** | **1.0e-04** | 6.1e-01 | **8.6e-03** | **6.8e-03** | **1.8e-03** | **1.8e-03** | **6.0e-04** | **1.0e-04** | **1.0e-04** | **1.0e-04** | **1.0e-04** |
| | | M2, M3 | **3.0e-02** | **1.0e-04** | 8.8e-01 | **4.2e-02** | **3.2e-02** | **4.5e-02** | **4.5e-02** | **2.6e-02** | **3.0e-02** | 8.8e-01 | **4.5e-02** | **2.6e-02** |
| | 5 kb | M1, M2 | **8.7e-03** | **2.5e-02** | 8.0e-01 | **3.6e-02** | **2.2e-02** | **5.0e-04** | **5.0e-04** | **2.0e-04** | **1.0e-04** | **1.0e-04** | **1.0e-04** | **1.0e-04** |
| | | M1, M3 | **4.0e-04** | **1.0e-04** | 9.1e-01 | **1.3e-03** | **5.0e-04** | **1.0e-04** | **1.0e-04** | **6.0e-04** | **1.0e-04** | **1.0e-04** | **1.0e-04** | **1.0e-04** |
| | | M2, M3 | **3.4e-02** | **1.0e-04** | 9.1e-01 | **4.8e-02** | **3.6e-02** | **5.0e-02** | **5.0e-02** | **3.7e-02** | **3.4e-02** | 9.1e-01 | **5.0e-02** | **3.7e-02** |
| | 10 kb | M1, M2 | **1.0e-04** | **3.0e-02** | 7.6e-01 | **1.0e-04** | **1.0e-04** | **1.0e-04** | **1.0e-04** | **1.0e-04** | **1.0e-04** | **1.0e-04** | **1.0e-04** | **1.0e-04** |
| | | M1, M3 | **1.0e-04** | **1.0e-04** | 3.3e-01 | **1.0e-04** | **1.0e-04** | **1.0e-04** | **1.0e-04** | **1.0e-04** | **1.0e-04** | **1.0e-04** | **1.0e-04** | **1.0e-04** |
| | | M2, M3 | **1.0e-02** | **1.0e-04** | 5.5e-01 | **1.2e-02** | **1.0e-02** | **7.9e-03** | **7.9e-03** | **7.1e-03** | **1.0e-02** | 5.5e-01 | **7.9e-03** | **7.1e-03** |
| | 15 kb | M1, M2 | **1.0e-04** | **3.5e-02** | 6.6e-01 | **1.0e-04** | **1.0e-04** | **1.0e-04** | **1.0e-04** | **2.0e-04** | **1.0e-04** | **1.0e-04** | **1.0e-04** | **1.0e-04** |
| | | M1, M3 | **1.0e-04** | **1.0e-04** | 4.5e-01 | **1.0e-04** | **1.0e-04** | **1.0e-04** | **1.0e-04** | **1.0e-04** | **1.0e-04** | **1.0e-04** | **1.0e-04** | **1.0e-04** |
| | | M2, M3 | **8.0e-04** | **1.0e-04** | 6.4e-01 | **2.0e-03** | **9.0e-04** | **9.0e-04** | **9.0e-04** | **5.0e-04** | **8.0e-04** | 6.4e-01 | **9.0e-04** | **5.0e-04** |
| resolution $\rho$ | 5 kb | M1, M2 | 7.8e-02 | 1.3e-01 | 1.2e-01 | 7.6e-02 | 7.8e-02 | **3.8e-03** | **3.8e-03** | **3.8e-03** | **1.0e-04** | **1.0e-04** | **1.0e-04** | **1.0e-04** |
| | | M1, M3 | **1.4e-02** | **1.2e-02** | 3.5e-01 | **1.2e-02** | **1.2e-02** | **3.0e-02** | **3.0e-02** | 6.2e-02 | **1.0e-04** | **1.0e-04** | **1.0e-04** | **1.0e-04** |
| | | M2, M3 | 4.7e-01 | 8.2e-02 | **9.9e-03** | 4.9e-01 | 4.7e-01 | 6.2e-01 | 6.2e-01 | 7.6e-01 | 4.7e-01 | **9.9e-03** | 6.2e-01 | 7.6e-01 |
| | 10 kb | M1, M2 | **5.6e-03** | **1.9e-02** | 7.4e-01 | **2.7e-03** | **4.0e-03** | **1.1e-03** | **1.1e-03** | **1.5e-03** | **1.0e-04** | **1.0e-04** | **1.0e-04** | **1.0e-04** |
| | | M1, M3 | **1.0e-04** | **1.0e-04** | **1.9e-02** | **1.0e-04** | **1.0e-04** | **2.0e-04** | **2.0e-04** | **2.0e-04** | **1.0e-04** | **1.0e-04** | **1.0e-04** | **1.0e-04** |
| | | M2, M3 | **4.9e-02** | **1.4e-02** | **2.0e-02** | 5.2e-02 | **4.9e-02** | 7.4e-02 | 7.4e-02 | 1.1e-01 | **4.9e-02** | **2.0e-02** | 7.4e-02 | 1.1e-01 |
| | 25 kb | M1, M2 | **1.0e-04** | **1.0e-04** | 6.3e-01 | **1.0e-04** | **1.0e-04** | **1.0e-04** | **1.0e-04** | **1.0e-04** | **1.0e-04** | **1.0e-04** | **1.0e-04** | **1.0e-04** |
| | | M1, M3 | **1.0e-04** | **1.0e-04** | **1.0e-04** | **1.0e-04** | **1.0e-04** | **1.0e-04** | **1.0e-04** | **1.0e-04** | **1.0e-04** | **1.0e-04** | **1.0e-04** | **1.0e-04** |
| | | M2, M3 | **1.0e-04** | **1.0e-04** | **1.0e-04** | **1.0e-04** | **1.0e-04** | **1.0e-04** | **1.0e-04** | **1.0e-04** | **1.0e-04** | **1.0e-04** | **1.0e-04** | **1.0e-04** |
| | 50 kb | M1, M2 | **1.0e-04** | **1.0e-04** | **2.0e-04** | **1.0e-04** | **1.0e-04** | **1.0e-04** | **1.0e-04** | **1.0e-04** | **1.0e-04** | **1.0e-04** | **1.0e-04** | **1.0e-04** |
| | | M1, M3 | **1.0e-04** | **1.0e-04** | **2.0e-04** | **1.0e-04** | **1.0e-04** | **1.0e-04** | **1.0e-04** | **1.0e-04** | **1.0e-04** | **1.0e-04** | **1.0e-04** | **1.0e-04** |
| | | M2, M3 | **1.0e-04** | **1.0e-04** | **5.0e-04** | **1.0e-04** | **1.0e-04** | **1.0e-04** | **1.0e-04** | **1.0e-04** | **1.0e-04** | **5.0e-04** | **1.0e-04** | **1.0e-04** |
| global | - | M1, M2 | **1.0e-04** | **1.0e-04** | 1.9e-01 | **1.0e-04** | **1.0e-04** | **1.0e-04** | **1.0e-04** | **1.0e-04** | **1.0e-04** | **1.0e-04** | **1.0e-04** | **1.0e-04** |
| | | M1, M3 | **1.0e-04** | **1.0e-04** | 1.8e-01 | **1.0e-04** | **1.0e-04** | **1.0e-04** | **1.0e-04** | **1.0e-04** | **1.0e-04** | **1.0e-04** | **1.0e-04** | **1.0e-04** |
| | | M2, M3 | **1.0e-04** | **1.0e-04** | 6.6e-01 | **1.0e-04** | **1.0e-04** | **1.0e-04** | **1.0e-04** | **1.0e-04** | **1.0e-04** | 6.6e-01 | **1.0e-04** | **1.0e-04** |

| cell line | accession number | measure | M1 | M2 | M3 | M4 |
|---|---|---|---|---|---|---|
| H1-hESC | 4DNFI9GMP2J8 | TPR | 14.13 | 11.48 | 9.71 | 3.84 |
| | | TNR | 93.90 | 97.77 | 97.97 | 99.24 |
| | | PPV | 35.65 | 55.14 | 53.30 | 54.78 |
| | | bACC | 54.02 | 54.62 | 53.84 | 51.54 |
| | | GM | 36.42 | 33.50 | 30.85 | 19.52 |
| | | F1c | 20.24 | 19.01 | 16.43 | 7.18 |
| | | FMc | 22.44 | 25.16 | 22.76 | 14.50 |
| | | JI | 11.26 | 10.50 | 8.95 | 3.72 |
| | | AHR | 59.90 | 11.48 | 9.71 | 17.00 |
| | | FGC | 79.08 | 54.92 | 53.65 | 99.12 |
| | | F1r | 68.17 | 18.99 | 16.45 | 29.02 |
| | | FMr | 68.83 | 25.11 | 22.83 | 41.05 |
| | | nAF | 32,644 | 6,257 | 5,294 | 9,263 |
| | | nSP | 26,210 | 11,765 | 12,018 | 3,864 |
| | 4DNFI6HDY7WZ | TPR | 13.84 | 7.61 | 5.07 | 2.50 |
| | | TNR | 92.19 | 98.31 | 98.99 | 99.40 |
| | | PPV | 29.76 | 51.86 | 54.60 | 49.89 |
| | | bACC | 53.01 | 52.96 | 52.03 | 50.95 |
| | | GM | 35.72 | 27.35 | 22.41 | 15.77 |
| | | F1c | 18.89 | 13.27 | 9.28 | 4.77 |
| | | FMc | 20.29 | 19.87 | 16.64 | 11.17 |
| | | JI | 10.43 | 7.11 | 4.87 | 2.44 |
| | | AHR | 60.29 | 7.61 | 5.07 | 11.83 |
| | | FGC | 71.89 | 51.48 | 54.60 | 98.11 |
| | | F1r | 65.58 | 13.26 | 9.28 | 21.11 |
| | | FMr | 65.84 | 19.79 | 16.64 | 34.07 |
| | | nAF | 32,856 | 4,147 | 2,765 | 6,446 |
| | | nSP | 31,496 | 8,415 | 5,989 | 2,757 |
| GM12878 | ENCFF993FGR | TPR | 17.76 | 2.57 | 4.70 | 4.08 |
| | | TNR | 96.33 | 97.74 | 96.56 | 99.33 |
| | | PPV | 41.23 | 14.15 | 16.54 | 47.01 |
| | | bACC | 57.05 | 50.15 | 50.63 | 51.71 |
| | | GM | 41.37 | 15.84 | 21.30 | 20.14 |
| | | F1c | 24.83 | 4.35 | 7.32 | 7.51 |
| | | FMc | 27.06 | 6.03 | 8.82 | 13.85 |
| | | JI | 14.18 | 2.22 | 3.80 | 3.90 |
| | | AHR | 60.32 | 2.57 | 4.70 | 15.55 |
| | | FGC | 82.70 | 14.17 | 16.56 | 97.54 |
| | | F1r | 69.76 | 4.35 | 7.32 | 26.83 |
| | | FMr | 70.63 | 6.03 | 8.82 | 38.95 |
| | | nAF | 21,970 | 935 | 1,712 | 5,665 |
| | | nSP | 18,127 | 6,762 | 14,076 | 3,173 |
| | ENCFF216QQM | TPR | 20.41 | 8.45 | 5.02 | 2.16 |
| | | TNR | 86.91 | 98.80 | 99.40 | 99.67 |
| | | PPV | 18.44 | 50.59 | 54.99 | 48.79 |
| | | bACC | 53.66 | 53.62 | 52.21 | 50.92 |
| | | GM | 42.11 | 28.89 | 22.34 | 14.68 |
| | | F1c | 19.37 | 14.48 | 9.20 | 4.14 |
| | | FMc | 19.40 | 20.67 | 16.62 | 10.27 |
| | | JI | 10.73 | 7.80 | 4.82 | 2.11 |
| | | AHR | 78.05 | 8.45 | 5.02 | 9.10 |
| | | FGC | 51.73 | 50.41 | 55.23 | 97.96 |
| | | F1r | 62.22 | 14.47 | 9.21 | 16.65 |
| | | FMr | 63.54 | 20.63 | 16.65 | 29.85 |
| | | nAF | 28,425 | 3,076 | 1,829 | 3,313 |
| | | nSP | 51,099 | 6,539 | 3,547 | 1,621 |

Supplementary Table 7. Classification and recognition measures for each real Hi-C matrix – resolution set to 10 kb. The following encoding is adopted: M1 = StripePy, M2=Chromosight, M3=StripeCaller, M4=Stripenn.

| tool | elapsed real time | max rss (Mb) | performance improcement |
|---|---|---|---|
| M1 | 1m 11.91s ± 0.18s | 4109.95 ± 8.54 | 2.13x |
| M2 | 4m 17.4s ± 0.41s | 2984.23 ± 30.85 | 7.61x |
| M3 | 33.82s ± 0.29s | 634.69 ± 0.53 | 1.0x |
| M4 | 37m 13.89s ± 6.44s | 1671.41 ± 12.33 | 66.06x |

Supplementary Table 8. Elapsed times, maximum resident set size, and performance improvements for contact map with accession number ENCFF993FGR – resolution set to 10 kb. The following encoding is adopted: M1 = StripePy, M2=Chromosight, M3=StripeCaller, M4=Stripenn.
Timings were measured on a workstation equipped with an AMD Ryzen 9 7950X3D 16-Core Processor, 4x32GB of DDR5 4800 MT/s RAM, and running Fedora 41 (Linux 6.12.11). Temporary and output files were written to an NVMe drive (PCIE 4.0) with plenty of free space available. Values shown in the table correspond to the average and standard error of measurements across 5 independent runs for each tool.

Supplementary Table 9. Comparing normalizations. We report classification and recognition measures, number of anchors found (nAF), and number of stripes predicted (nSP) obtained when running StripePy on the real Hi-C contact maps with no (default), ice and scale normalizations. Here, resolution is set to 10 kb.

| | GM12878 | | | | | | H1-hESC | | | | | |
|---|---|---|---|---|---|---|---|---|---|---|---|---|
| | 4DNFI9GMP2J8 | | | 4DNFI6HDY7WZ | | | ENCFF993FGR | | | ENCFF216QQM | | |
| measure | NONE | ICE | SCALE | NONE | ICE | SCALE | NONE | ICE | SCALE | NONE | ICE | SCALE |
| TPR | 14.13 | 4.24 | 7.44 | 13.84 | 7.09 | 7.57 | 17.76 | 7.78 | 11.10 | 20.41 | 10.26 | 12.15 |
| TNR | 93.90 | 98.28 | 98.00 | 92.19 | 96.92 | 96.87 | 96.33 | 98.08 | 97.96 | 86.91 | 95.95 | 95.00 |
| PPV | 35.65 | 37.05 | 47.03 | 29.76 | 35.48 | 36.67 | 41.23 | 37.04 | 44.06 | 18.44 | 26.88 | 26.06 |
| bACC | 54.02 | 51.26 | 52.72 | 53.01 | 52.00 | 52.22 | 57.05 | 52.93 | 54.53 | 53.66 | 53.10 | 53.58 |
| GM | 36.42 | 20.42 | 27.01 | 35.72 | 26.20 | 27.09 | 41.37 | 27.62 | 32.97 | 42.11 | 31.37 | 33.97 |
| F1c | 20.24 | 7.61 | 12.85 | 18.89 | 11.81 | 12.55 | 24.83 | 12.86 | 17.73 | 19.37 | 14.85 | 16.57 |
| FMc | 22.44 | 12.54 | 18.71 | 20.29 | 15.86 | 16.66 | 27.06 | 16.97 | 22.11 | 19.40 | 16.60 | 17.80 |
| JI | 11.26 | 3.96 | 6.87 | 10.43 | 6.28 | 6.70 | 14.18 | 6.87 | 9.73 | 10.73 | 8.02 | 9.04 |
| AHR | 59.90 | 27.02 | 34.33 | 60.29 | 35.58 | 37.43 | 60.32 | 35.96 | 42.21 | 78.05 | 45.87 | 51.00 |
| FGC | 79.08 | 92.69 | 90.08 | 71.89 | 81.06 | 83.11 | 82.70 | 87.66 | 87.15 | 51.73 | 67.75 | 64.28 |
| F1r | 68.17 | 41.84 | 49.71 | 65.58 | 49.46 | 51.62 | 69.76 | 51.00 | 56.87 | 62.22 | 54.70 | 56.87 |
| FMr | 68.83 | 50.04 | 55.61 | 65.84 | 53.71 | 55.77 | 70.63 | 56.15 | 60.65 | 63.54 | 55.74 | 57.26 |
| nAF | 32,644 | 14,722 | 18,708 | 32,856 | 19,390 | 20,398 | 21,970 | 13,098 | 15,373 | 28,425 | 16,705 | 18,575 |
| nSP | 26,210 | 6,303 | 8,819 | 31,496 | 11,671 | 11,805 | 18,127 | 7,753 | 9,335 | 51,099 | 14,210 | 17,498 |
